# Neo-functionalization and co-option of Pif genes facilitate the evolution of a novel shell microstructure in oysters

**DOI:** 10.1101/2023.05.22.541698

**Authors:** Yitian Bai, Shikai Liu, Yiming Hu, Hong Yu, Lingfeng Kong, Chengxun Xu, Qi Li

## Abstract

Molluscan shell, composed of a diverse range of architectures and microstructures, is a classic model system to study the relationships between molecular evolution and biomineralized structure formation. The shells of oysters differ from those of other molluscs by possessing a novel microstructure, chalky calcite, which facilitates adaptation to the sessile lifestyle. However, the genetic basis and evolutionary origin of this adaptive innovation remain largely unknown. Here, we present the first chromosome-level genome and shell proteomes of the Iwagaki oyster *Crassostrea nippona*. Multi-omic integrative analyses revealed that independently evolved and co-opted genes as well as lineage-specific domains are involved in the formation of chalky layer in the oysters. Rapid mineralization involving chalky calcite are essential for reconstruction of the shell. Importantly, von Willebrand factor type A and chitin-binding domains are identified as basic members of molluscan biomineralization toolkit. We show that the well-known Pif shared a common origin in the last common ancestor of Bilateria. Furthermore, Pif and LamG3 genes acquire new genetic function for shell mineralization in bivalves and the chalky layer formation in oysters through a combination of gene duplication and domain reorganization. Our findings highlight neo-functionalization as a crucial mechanism for shell diversity, which may be applied more widely for studies on the evolution of metazoan biomineralization. This study also has potential implications for material science and biomimetic research.

## Introduction

Biomineralized exoskeleton represents a key evolutionary innovation that contributes to the rapid diversification of living organisms dating back to the early Cambrian (Knoll 2003). Among mineralizing metazoans, Mollusca particularly benefits from the various functions of mineralized shell (McDougall and Degnan 2018), resulting in the evolutionary and ecological success of this extremely diverse phylum (Wanninger and Wollesen 2019). Molluscan shells are composed of calcium carbonate crystals and multiple organic matrix components comprising proteins, polysaccharides, and lipids (Agbaje, et al. 2018). In spite of a minor organic part in shell by mass, shell matrix proteins (SMPs) play the critical roles in shell construction (Clark, et al. 2020). Over the past decades, a large array of SMPs have been identified from molluscan shells (Marin 2020). These proteins are not only involved in the formation of organic framework, but also control the nucleation and growth of calcium carbonate polymorphs (calcite or aragonite) (Falini, et al. 1996; Addadi, et al. 2006; Ponce and Evans 2011).

Generally, molluscan shells contain multiple layers, and each layer is characterized by a specific shell microstructure (e.g., prismatic, crossed-lamellar, nacreous, and homogeneous) (Chateigner, et al. 2000). Although shell morphology is conserved within many taxa, ultrastructural and molecular analyses revealed that shell microstructures have evolved independently multiple times in many molluscan lineages (Bieler, et al. 2014; McDougall and Degnan 2018). At the proteomic level, distinct partitioning of SMPs were observed between prismatic and nacreous layers (Marie, et al. 2012; Shimizu, Negishi, et al. 2022). Molecular innovation promotes the evolution of multiple novelties in emerging lineages (Kaessmann 2010; Erwin 2015). Likewise, the diversity of shell microstructures is associated with the molecular evolution of genes acting as SMPs (Kocot, et al. 2016; Aguilera, et al. 2017). Novel genes encoding SMPs are often formed by gene duplication and combinations of recruited/co-opted protein domains from ancient genes (Kocot, et al. 2016). Such SMPs usually undergo rapid functional evolution (neo-functionalization or sub-functionalization) (Shimizu, Takeuchi, et al. 2022). Biomineralization evolved independently but convergently across phyla (Gilbert, et al. 2022), implying that diverse biomineralized structures of metazoans may share common molecular evolutionary dynamics that transcends biological differences. Consequently, the genetic dissection of the molecular relationship between SMPs and shell microstructures provide an excellent example for understanding the origin and functional evolution of novel genes, especially involved in metazoan biomineralization.

The bivalve family Ostreidae, also called oyster, is the only one molluscan group that possesses chalky calcite as a shell layer (Dauphin, et al. 2013). A previous study identified an EGF-like domain-containing (ELC) protein in the chalky layer of *Crassostrea gigas*, and revealed that this protein participated in crystal aggregation during shell mineralization (Iwamoto, et al. 2020). However, the molecular details of chalky layer formation remain enigmatic. Even more elusive is the evolutionary origin of the novel shell microstructure. In this study, we focus on a member of Ostreidae family, the Iwagaki oyster *C. nippona*. As an ideal model for studying shell mineralization, *C. nippona* has considerable chalky calcite deposited in its left shell. To better understand the genetic basis of chalky layer formation, we assembled a chromosome-level genome of *C. nippona*, and generated comprehensive transcriptomic resources. In addition, proteomic analyses of three types of *C. nippona* shell layers (prismatic, foliated, and chalky layers) were constructed using liquid chromatography tandem mass spectrometry (LC-MS/MS). Together, these data provide a molecular explanation for the evolution of shell microstructures, and highlights the origin and functional evolution of key genes underlying the diversity of molluscan shell mineralization.

## Results and discussion

### Genome assembly and annotation

A chromosome-scale genome assembly of *C. nippona* was constructed from 27.5 Gb (∼ 67-fold coverage) of PacBio high-fidelity circular consensus sequencing (HiFi-CCS) reads and 60.8 Gb of Hi-C sequence data (fig. 1a; supplementary fig. S1a). The assembly has a total length of 530.1 Mb (Scaffold N50 = 50.9 Mb) and a GC content of 33.8% (supplementary table S1). The assembled genome size was consistent with the estimation based on k-mer analysis (k = 17) (supplementary fig. S1b), and the result from flow cytometry analysis in a previous study (Adachi, et al. 2021). The high quality of our genome assembly was supported by over 99% mapping rate of sequencing reads (supplementary table S1) and 97.3% of Benchmarking Universal Single-Copy Orthologs (BUSCO) completeness against the metazoan core gene set (supplementary table S2).

**Fig. 1.**
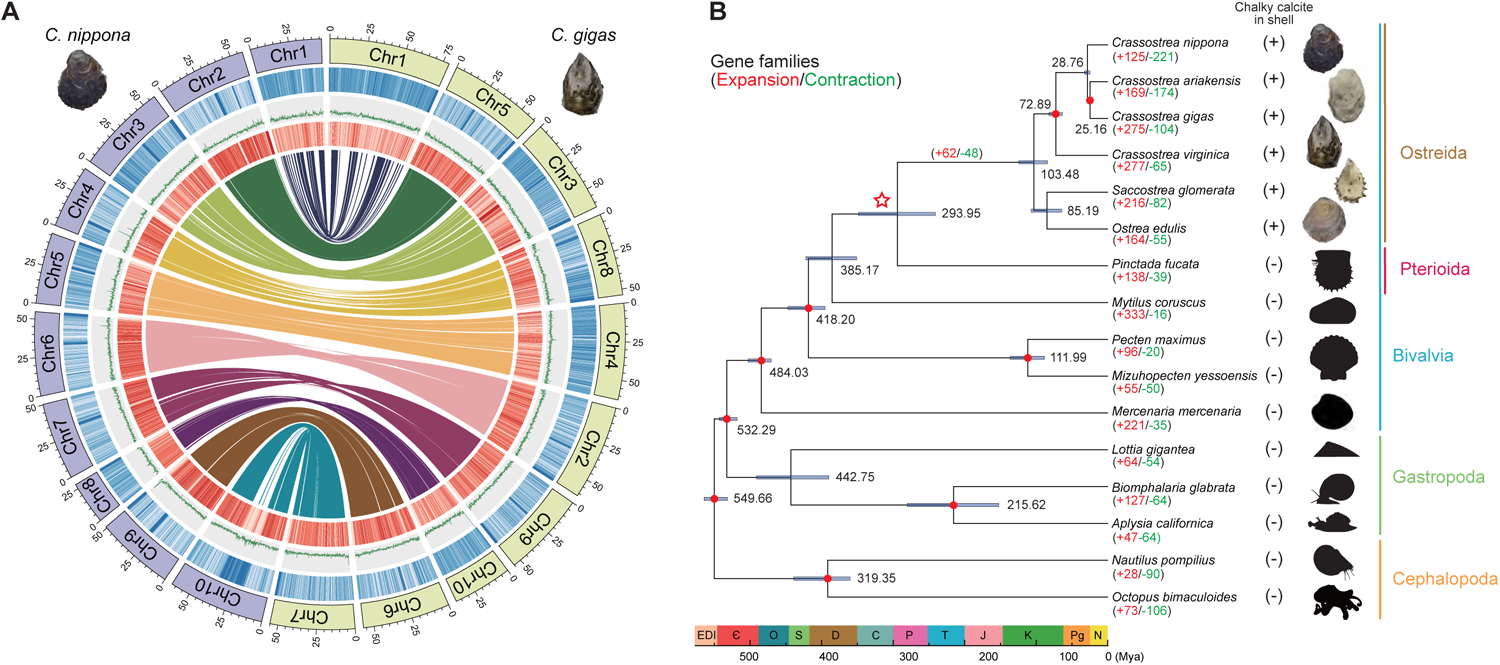
Genomic landscapes of two oysters and the origin of chalky layer in molluscs. **(A)** Circos plots showing conserved synteny between *Crassostrea nippona* (left) and *C. gigas* (GCA_011032805.1) (right). From outer to inner circle: repeat coverage, GC content, gene distribution, genomic synteny. The sliding window size is 100 kb. **(B)** Phylogenetic tree of 16 representative molluscs. Nine nodes with red dots represent reference-calibrated time points (supplementary table S7). The node with a red hollow star shows the divergence time between Ostreida and Pterioida. The plus signs after species names indicate chalky calcite in the shell; minus signs mean no chalky calcite in the shell. The purple horizontal lines indicate the 95% confidence intervals of divergence times. Numbers of gene families undergoing expansion and contraction for each lineage are shown in red and green, respectively. The color labelling scheme of taxa: Bivalvia: blue; Gastropoda: green; Cephalopoda: orange; Ostreida: brown; Pterioida: purplish red. Abbreviations: Mya, million years ago; Є, Cambrian; C, Carboniferous; D, Devonian; EDI, Ediacaran; J, Jurassic; K, Cretaceous; N, Neogene; O, Ordovician; P, Permian; Pg, Paleogene; S, Silurian; T, Triassic.

A large portion (44.69%) of the *C. nippona* genome was annotated as repetitive elements, which were dominated by DNA transposons (17.90%) and Helitrons (13.35%) (supplementary fig. S2 and table S3). In total, 28,871 protein coding genes were predicted in the *C. nippona* genome, with an average gene length of 9005 bp (supplementary table S4). Approximately 96% of predicted genes were functionally annotated using various public databases (supplementary table S5). Assessment of the predicted genes using BUSCO analysis revealed that 94.8% of the predicted genes were complete ortholog genes (supplementary table S2). In addition, a total of 5401 non-coding RNA (ncRNA) genes were identified in the *C. nippona* genome, including 2474 tRNAs, 1847 rRNAs, 1036 snRNAs, and 44 miRNAs (supplementary table S6).

### Genomic rearrangements in oysters

We found a high level of one-to-one syntenic relationship at the chromosome level in *Crassostrea* species (fig.1a, supplementary fig. S3a), which was consistent with observations reported in previous studies (Li, Dai, et al. 2021; Zhang, et al. 2022; Dong, et al. 2023; Li, et al. 2023). Moreover, various inter-chromosomal rearrangements were observed between *C. nippona* and *Mizuhopecten yessoensis* (supplementary fig. S3b). The *M. yessoensis* was reported to possess a highly conserved 19-chromosome karyotype similar to that of the bilaterian ancestor (Wang, et al. 2017). The 10-chromosome karyotype of *C. nippona* was mainly derived from fusions of two or more ancestral chromosome segments of *M. yessoensis*, except for Chr1, which was only originated from an ancestral chromosome (supplementary fig. S3b). Similar chromosome rearrangements were also observed between *C. nippona* and *Pinctada fucata* (supplementary fig. S3b). Together, a combination of fission, fusion, and retention from the ancestral chromosomes resulted in the 10 chromosomes of oysters. Chromosomal fusions and fissions were found to facilitate adaptation to divergent environments by creating regions of low recombination that enabled the formation of adaptative clusters (Liu, et al. 2022). Therefore, we speculate that large-scale fission and fusion rearrangements from ancestral chromosomes may contribute to the genomic adaptation of oysters to sessile life in a highly stressful environment (Zhang, et al. 2012).

### Comparative genomics reveal the origin of chalky calcite

We selected 15 molluscan species with whole genome assembly for comparative genomic analyses with *C. nippona* (supplementary table S2). A total of 1253 one-to-one single-copy orthologous genes were identified and used for the construction of phylogenetic tree (fig. 1b, supplementary table S7). Molecular clock analysis based on the secondary calibrations suggested that the common ancestor of *C. nippona* diverged from *C. gigas* and *C. ariakensis* at 28.76 million years ago (Mya) (25.37-32.76 Mya) (fig. 1b), in agreement with evidence from a previous study (Li, Kou, et al. 2021). Within Bivalvia, the Ostreida and Pterioida diverged at 293.95 Mya (241.02-348.46 Mya) (fig. 1b). Given the origin and the early divergence of Ostreida species (Guo, et al. 2018; Li, Kou, et al. 2021), our result supports the hypothesis that the Permian was a key period for the radiation of bivalves, and molluscan ecological dominance first occurred prior to the end-Permian mass extinction (Clapham and Bottjer 2007). Notably, we found that the Ostreidae-specific chalky calcite in the oyster shells occurred after the divergence of the Ostreida and Pterioida (fig. 1b). This novel shell microstructure is a result of independent evolution in oysters. Global environmental change at the end of the Permian Period disturbed the diversity of biomineralization (Gilbert, et al. 2022). The common ancestor of Ostreida species may have to respond in distinct ways (e.g., chalky deposition in the shell) to selective pressure at the end-Permian mass extinction.

Among the selected genomes, we identified a set of 620 Ostreidae-specific gene families and 62 expanded gene families, respectively. Gene Ontology (GO) enrichment analyses of these gene families revealed the components involved in shell mineralization of oysters such as cell adhesion, extracellular matrix, fibronectin binding, and chitin binding (supplementary table S8 and S9). Moreover, gene families known to produce proteins involved in shell formation, such as tyrosinase (Feng, et al. 2019), peroxidase (Chen, et al. 2019), and tissue inhibitor of metalloproteinase (TIMP) (Yan, et al. 2014), have undergone large independent expansions in Ostreidae (supplementary figs. S4a, S5a and S6a). The majority of these gene members were highly expressed in the mantle of *C. nippona* (supplementary figs. S4b, S5b, S6b, and S7). Combined with shell proteomes of *C. nippona* (supplementary table S10), some of oyster-specific gene members were classified as SMPs in the chalky layer (supplementary figs. S4, S5 and S6), suggesting their co-option into the evolution of the chalky calcite in the oyster shells.

### Evolution and formation of chalky calcite in oysters

Like other oyster species (Checa, et al. 2009; Dauphin, et al. 2013; Fitzer, et al. 2019; Clark, et al. 2020), *C. nippona* has a pure calcite shell, with an outer layer of calcite prisms and inner multi-layered structures consisting of repeated foliated and chalky layers (fig. 2a, supplementary fig. S8). In the multi-layer, the foliated layer is stacked by dense sheets of folia, while chalky structures are constituted of loose calcite blades with ample interconnected porosity (fig. 2a). Foliated calcite is a common microstructure in many pteriomorphian bivalves (Lemer, et al. 2016), whereas chalk is found exclusively in oysters (Dauphin, et al. 2013). The diversity of shell microstructure is usually associated with life-habits (Lemer, et al. 2016). Oyster has a sessile lifestyle supported by cemented attachment. As a filler with remarkable plasticity, chalky deposition allows oysters to closely cement their left shell with uneven substrates in estuarine or intertidal zones (Banker and Sumner 2020). Chalky microstructure also enhances the crack resistance of the oyster shell, which may serve a similar function to the holes in bones (MacDonald, et al. 2009), implying the potential convergent evolution of biomineralized skeletons in oysters and vertebrates. In addition, compared with that of right shell, chalky deposition is more frequent in the left valve of oyster shell (Checa, et al. 2018), which may act as a driver of left-right shell asymmetry in oysters. Thus, the chalky layer may result from adaptive evolution for the sessile life of oysters.

**Fig. 2.**
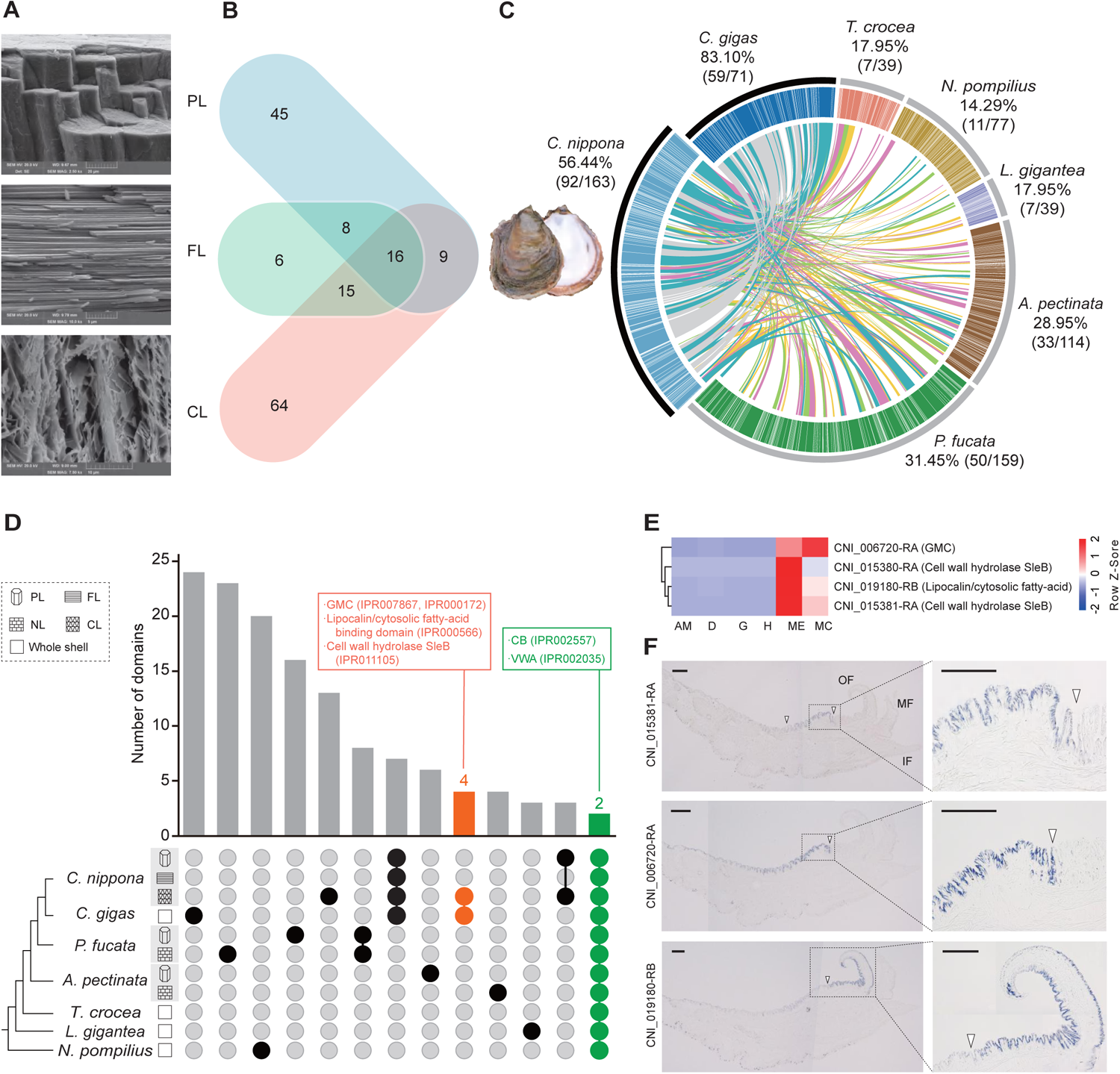
Microstructures and proteomes of the *Crassotrea nippona* shell. Abbreviations: PL, prismatic layer; FL, foliated layer; CL, chalky layer; NL, nacre layer. **(A)** SEM micrographs of the microstructures of the prismatic, foliated, and chalky layers of the *C. nippona* shell. **(B)** Number of proteins identified from the prismatic, foliated, and chalky layers. **(C)** Circos diagram of seven representative molluscan shell proteomes (the E-value cut-off of BLASTP is 1e-20); with the shell of farmed *C. nippona* on the left. Protein pairs sharing sequence similarity in the top quartile are linked in blue, the second quartile is linked in pink, third quartile has green links and lowest quartile of similarity has yellow links. Oyster-specific protein pairs are linked by grey lines. Percentages and proportions in brackets indicate the number of SMPs having similarities between *C. nippona* and the other six molluscs. **(D)** Upset plot comparing the protein-domains identified from the shell proteomes of seven molluscs. Domains only detected in the *C. gigas* shell and the chalky layer of *C. nippona* shell are indicated in orange. Domains shared by all shell layers across seven molluscs are colored in green. The complete results are shown in supplementary fig. S12. **(E)** Expression of SMPs with domains specifically detected in *C. gigas* and the chalky layer of *C. nippona* in six tissues of *C. nippona*. Abbreviations: AM, adductor muscle; D, digestive gland; G, gill; H, hemolymph; ME, mantle edge; MC, central mantle. **(F)** Spatial expression patterns of three SMPs with domains specific for chalky layers in the mantle tissue of *C. nippona*. Full view (left) and partial enlargement (right) show positive cells stained in blue by in situ hybridization of each gene, respectively. White arrows symbolize the end of the gene expression region. Scale bar, 200 µm. Abbreviations: OF, outer fold; MF, middle fold; IF, inner fold.

The formation of chalky microstructure was associated with an amino acid site mutation of an ELC protein by allowing the cleaved N-terminal region of ELC protein to be incorporated into the chalky layer of *C. gigas* (Iwamoto, et al. 2020). However, shell matrix proteins (SMPs) are elements of a comprehensive regulatory network and work cooperatively to form a given microstructure (Marin 2020). The production of shell microstructure cannot be solely attributed to certain separated constituents. To comprehensively investigate the molecular basis of chalky layer formation, we performed proteome sequencing of the prismatic, foliated, and chalky layers from the *C. nippona* shell and identified a total of 78, 45, and 104 SMPs, respectively (fig. 2b, supplementary fig. S9, table S10). SMPs are secreted by the epithelial cells on the dorsal region of mantle. Expression patterns of genes encoding SMPs showed that most of these genes (82.2%) were highly expressed in the mantle (supplementary fig. S10), confirming the essential role of the mantle in shell mineralization as previously reported (Marie, et al. 2012; Zhang, et al. 2012; Zhao, et al. 2018; Zhang, et al. 2021). Comparative shell proteomic analysis indicated that 92 of *C. nippona* SMPs (56.4%) shared similarity with sequences in shell proteomes of the other six molluscs, including four bivalves, one gastropod, and one cephalopod (fig. 2c). A high degree of unique matches was found between *C. nippona* and *C. gigas*, and the least number of matches was observed between *C. nippona* and *Nautilus pompilius* (fig. 2c). The results are not only consistent with evolutionary divergence times, but also attributed to the crystal polymorphs of the shells of different species, as the shells of *C. nippona* and *C. gigas* are entirely calcite (Dauphin, et al. 2013), whereas the *N. pompilius* shell is completely aragonite (Zhang, et al. 2021). The phylogenetic orthology analysis revealed 43 *C. nippona*-specific SMPs (supplementary fig. S11a). Interestingly, 26 of these SMPs were identified from the chalky layer (supplementary fig. S11b), thereby allowing us to infer that most of the SMPs involved in the formation of chalky layer have evolved independently.

Further protein domain analysis was performed across SMPs from different shell microstructures of seven molluscs (fig. 2d). Three domains were observed in both the *C. gigas* shell and the chalky layer of the *C. nippona* shell, including glucose-methanol-choline (GMC) oxidoreductase, lipocalin/cytosolic fatty-acid binding, and cell wall hydrolase SleB (fig. 2d). These domains represented a unique repertoire occurred exclusively in the chalky layer. SMPs containing the unique domains were highly expressed in the outer mantle epithelium of *C. nippona* (figs. 2e and 2f), confirming their participation in the formation of chalky layer. Notably, a previous study identified the GMC oxidoreductase domain unique to *C. gigas* by comparing shell proteomes of bivalves (Arivalagan, et al. 2017). The GMC domain was discussed as the roles in the development, immunity, and chemical defense of insects (Iida, et al. 2007; Sun, et al. 2012; Rahfeld, et al. 2014). However, the molecular function of GMC domain has remained largely unknown in molluscs. Similarly, the actual roles of lipocalin/cytosolic fatty-acid binding domain and cell wall hydrolase SleB domain still require further characterization in molluscan species.

### Rapid shell reconstruction with chalky calcite

Molluscs have the ability to repair damaged shells caused by external aggressors (Fleury, et al. 2008). Shell repair in oyster appeared to proceed faster than that of other bivalves (Yarra, et al. 2021). We compared gene expression patterns in mantle edge (ME) and central mantle (MC) from the shell-drilled *C. nippona* and non-drilled individuals, respectively (fig. 3, supplementary fig. S13, tables S11 and S12). Notably, most of differentially expressed genes (DEGs) encoding SMPs were up-regulated in the ME of the non-drill oysters (fig. 3a, supplementary table S11), whereas DEGs encoding SMPs were highly expressed in the MC during shell repair (fig. 3b, supplementary table S12). Under normal conditions, the physiological functions of ME and MC are shell growth and maintenance, respectively. However, shell damage may result in MC switching from maintenance to active repair and reconstruction (Yarra, et al. 2021). In oyster shell, chalky calcite is irregularly distributed in the inner layers (Dauphin, et al. 2013; Checa, et al. 2018), and acts as a cheap structural material to facilitate rapid growth of shell (Banker and Sumner 2020). During shell repair in *C. nippona*, plentiful chalky calcite was deposited following the foliated layer (figs. 3c and d), and many DEGs encoding SMPs of chalky layer were up-regulated in MC (fig. 3b, supplementary table S12). Therefore, the chalky deposition may promote the rapid reconstruction of oyster shell, thereby facilitating the environmental adaptation of oysters.

**Fig. 3.**
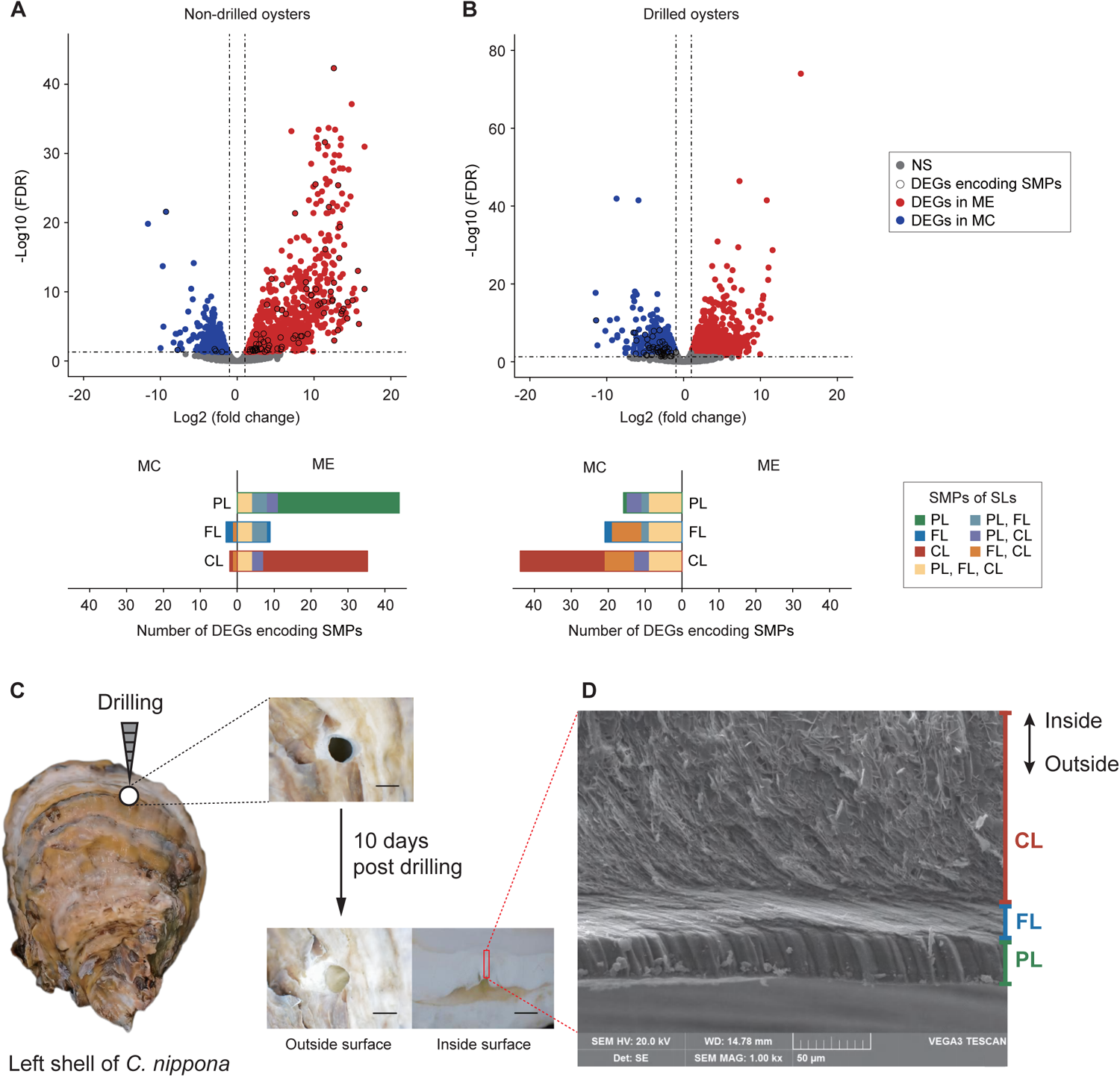
Rapid reconstruction of the *Crassostrea nippona* shell. Abbreviations as in fig 2, and DEGs, differentially expressed genes; ME, mantle edge; MC, mantle center; NS, Non-significant expression; SLs, shell layers. **(A)** Differential gene expression and number of DEGs encoding SMPs in mantle tissues of non-drilled *C. nippona* and **(B)** drilled oysters. **(C)** Schematic illustration of shell-drilling experiment and observation of shell regeneration process of *C. nippona*. Scale bar, 5 mm. **(D)** Ultrastructure of cross section of the whole repaired shell.

### Conversed CB and VWA domains in diverse shell microstructures

A biomineralization toolkit containing conserved domains within the Bivalvia was well-studied using shell proteomes or mantle transcriptomes (Arivalagan, et al. 2017; Yarra, et al. 2021). Although it was unclear about the detailed functions of these domains, the toolkit represented the core requirements for shell biomineralization. In our study, only two domains were completely conserved in the SMPs from different shell microstructures across molluscs (fig. 2d). These domains were chitin-binding (CB) and von Willebrand factor type A (VWA), which were also classified as the members of biomineralization toolkit in previous studies (Arivalagan, et al. 2017; Zhao, et al. 2018; Yarra, et al. 2021; Zhang, et al. 2021; Cavallo, et al. 2022; Shimizu, Negishi, et al. 2022). Based on these two domains, various SMPs were clustered into an orthogroup (OG0000000) which was shared by the shell proteomes of seven molluscs (supplementary fig. S11a and table S13). These evidences point to the possibility that CB and VWA domains are essential for shell formation, and represent the ancestral and basic components of molluscan biomineralization toolkit.

CB and VWA domains are widely distributed in diverse metazoan lineages (fig. 4). The VWA domain is often found in extracellular matrix proteins, and has an adhesion function through protein-protein interaction (Whittaker and Hynes 2002). The CB domain exhibits the high binding affinity to chitin, and plays a critical role in the construction of various biomineralized exoskeletons (Luo, et al. 2015; Jin, et al. 2019; Zhang, et al. 2019; Sun, et al. 2020; Sun, et al. 2021). Chitin is one of the major polysaccharides comprising the calcified shells of molluscs (Falini, et al. 1996). A chitinous scaffold provides the basic framework for interactions between extracellular matrix and calcium carbonates (Nudelman 2015; Du, et al. 2017). CB and VWA domains have both been found to participate in chitin-scaffolding and arranging calcium carbonate crystals of molluscan shell (Du, et al. 2017; Jin, et al. 2019). Thus, the combinations of VWA and CB domains may be associated with the diversity of shell microstructures.

**Fig. 4.**
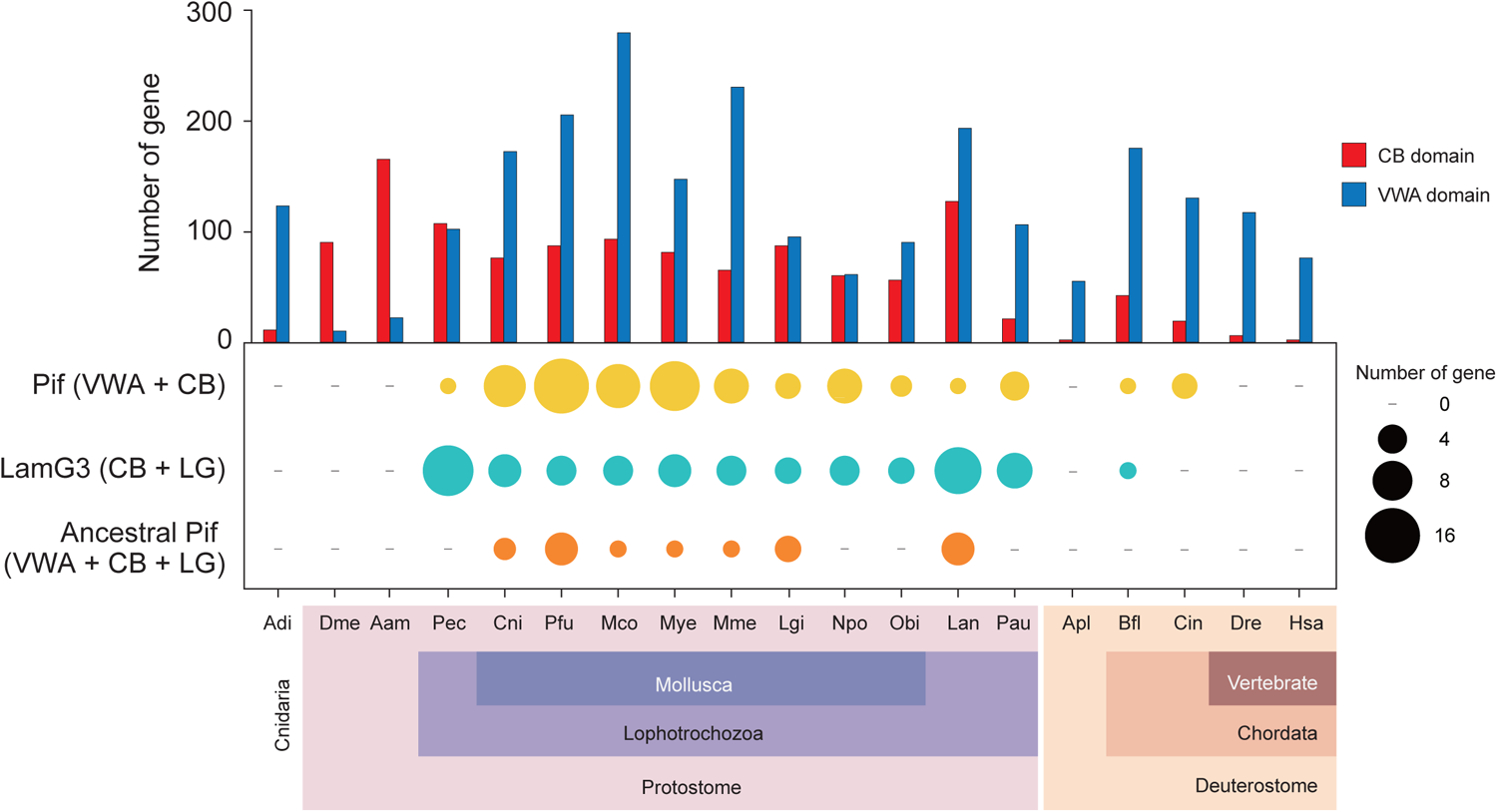
Distribution of chitin-binding (CB) and von Willebrand factor type A (VWA) domain containing genes among metazoans. Numbers of CB and VWA domain-containing genes (histogram graph) as well as Pif, ancestral Pif, and LamG3 genes (bubble chart) were determined in 19 metazoan genomes. Abbreviations: Aam, *Amphibalanus amphitrite*; Adi, *Acropora digitifera*; Apl, *Acanthaster planci*; Bfl, *Branchiostoma floridae*; Cin, *Ciona intestinalis*; Cni, *Crassostrea nippona*; Dme, *Drosophila melanogaster*; Dre, *Danio rerio*; Hsa, *Homo sapiens*; Lan, *Lingula anatina*; Lgi, *Lottia gigantea*; Mco, *Mytilus coruscus*; Mye, *Mizuhopecten yessoensis*; Mme, *Mercenaria mercenaria*; Npo, *Nautilus pompilius*; Obi, *Octopus bimaculoides*; Pau; *Phoronis australis*; Pec, *Paraescarpia echinospica*; Pfu, *Pinctada fucata*.

### Evolution of Pif proteins

The well-known combination of VWA and CB domains is Pif protein (Suzuki, et al. 2009). This acidic matrix protein and its homologs were not only identified in molluscan shells (Marie, et al. 2012; Suzuki, et al. 2013; Zhao, et al. 2018), but also involved in the construction of other mineralized structures of molluscs, such as sclerites and shell-like eggcase (Varney, et al. 2021; Yoshida, et al. 2022). Despite VWA and CB domains are common across metazoans, we found that Pif proteins were exclusively present in lophotrochozoans and two chordates (*Branchiostoma floridae* and *Ciona intestinalis*) (fig. 4). The conserved domain architecture with VWA, CB, and concanavalin A-like lectin/glucanase (LG) domains was previously considered as an ancestral Pif, which occurred in the last common ancestor (LCA) of Mollusca and Brachiopoda (Suzuki, et al. 2013; Zhao, et al. 2018). While in our study, the ancestral Pif proteins was not only observed in the genomes of bivalves, gastropods, and the brachiopod *Lingula anatine*, but also in other lophotrochozoans and chordates (fig. 4). Interestingly, Pif and its homologs, LamG3 proteins, were uncovered in the SMPs of *C. nippona* (supplementary fig. S15). The LamG3 proteins, composed of both CB and LG domains yet no VWA domain, were present in lophotrochozoans and *B. floridae* (fig. 4). Taken together, LamG3, Pif, and ancestral Pif shared similar structures and distribution in metazoans.

To further understand the evolutionary relationships of Pif and LamG3, a molecular phylogenetic tree was constructed using Pif, ancestral Pif, and LamG3 proteins from 16 metazoans (fig. 5a, supplementary table S14). Interestingly, one Pif from *M. yessoensis*, an ancestral Pif of *C. intestinalis*, and a LamG3 in *Lottia gigantea* were grouped into a single clade, supporting the intimate evolutionary relationship of Pif, ancestral Pif, and LamG3 in molluscs and chordates. The molluscan Pif and LamG3 genes formed separate monophyletic groups from other lophotrochozoans and chordates, suggesting that their evolutionary origin is prior to the split of deuterostomes and protostomes. Many of these genes were highly expressed in the mantle of molluscs (fig. 5a). Particularly, three groups of these genes were all highly expressed in the mantle tissues of bivalves (fig. 5a), and three members were expressed in dorsal region of the outer epithelium of the mantle of *C. nippona* (fig. 5b). These results implied that the three groups of Pif and LamG3 genes have potential functions in shell mineralization. However, a number of Pif did not exhibit high expression level in shell-forming mantle tissues of molluscs (fig. 5a). In addition, no Pif proteins were found in the shell proteomes of brachiopods (Immel et al. 2015; Isowa et al. 2015; Luo et al. 2015) and tube proteome of *Paraescarpia echinospica* (Sun, et al. 2021). Our findings are consistent with the hypothesis that the ancestral function of Pif may be not related to biomineralization (Zhao et al., 2018). Thus, Pif and LamG3 were independently co-opted for shell mineralization in the molluscan lineage.

**Fig. 5.**
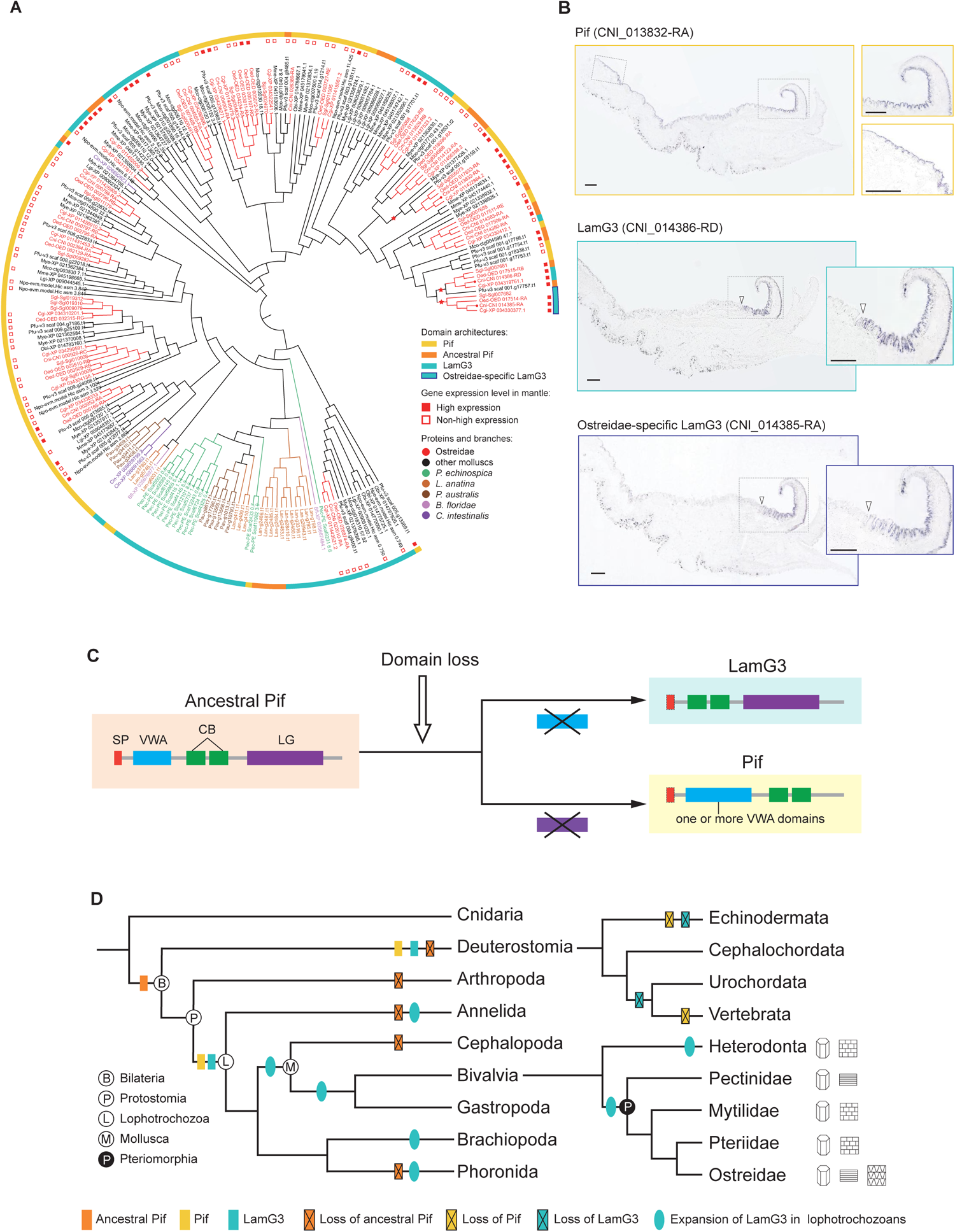
Evolution of Pif and LamG3 in metazoans. **(A)** Maximum likelihood tree of Pif and LamG3 in selected metazoans. The different colors of proteins and branches represent distinct lineages. Red solid squares indicate the genes highly expressed in the mantle of selected molluscan species, while the red hollow squares indicated genes that were not highly expressed in the mantle. Red stars on the clades represent three groups of genes which are all highly expressed in the mantles. **(B)** Spatial expression patterns of Pif and LamG3 in *C. nippona* mantle. The genes used for in situ hybridization were indicated with red dots in Fig. 5A. Scale bar, 200 µm. White arrows symbolize the end of the gene expression region. **(C)** The evolutionary model of LamG3, Pif, and ancestral Pif. Dash box indicate potential presence of SP domain. Abbreviations: SP, signal peptide; CB, chitin-binding domain; VWA, von Willebrand factor type A; LG, concanavalin A-like lectin/glucanase domain. **(D)** Reconstructions of evolution of Pif and LamG3 in metazoans. Microstructure models of shell layers as in fig 2d.

We further proposed an evolutionary model of Pif, ancestral Pif, and LamG3 (fig. 5c). Briefly, the ancestral Pif underwent the domain loss of VWA and LG, resulting in generation of LamG3 and Pif, respectively. The molecular phylogenetic analysis indicated that Pif and LamG3 had a common origin in Bilateria (fig. 5a and 5d). However, different major clades may diverge from multiple copies of the ancestral Pif in the LCA of Bilateria (fig. 5a). Although our analysis was based on the genomes of a limited number of species, the results indicated that the Pif and LamG3 genes in Lophotrochozoa and Deuterostomia convergently evolved from the ancestral Pif, then were lost in Echinodermata and Vertebrata (fig. 5d). In addition, the ancestral Pif gene was preserved in Bivalvia, Gastropoda, and Brachiopoda, but lost in other lophotrochozoans as well as Arthropoda and Deuterostomia. Notably, LamG3 gene has expanded at least once in many lophotrochozoan lineages (fig. 5d). These expansions may be linked to the diversity of biomineralization in molluscs and other lophotrochozoans (Sun, et al. 2021).

### Pif and LamG3 undergo neo-functionalization

Strikingly, an Ostreidae-specific LamG3 was identified as the SMP in the chalky layer of *C. nippona* (figs. 5a, supplementary fig. S15). Deep mining of the oyster genomes revealed that the Ostreidae-specific LamG3, as well as a set of LamG3 and Pif genes, were clustered into a clade (figs. 5a) and localized across the same chromosome (supplementary fig. S16). Therefore, we concluded with a hypothesized evolutionary process of origin and functionalization of the Pif_LamG3_cluster in bivalves (fig. 6). This gene cluster was evolved from a single copy of ancestral Pif gene (Pif-a) in the LCA of Mollusca. Firstly, two successive reverse tandem duplication of the ancestral Pif gene occurred in Bivalvia, and the region encoding the C-terminal VWA domain in the Pif-a gene was deleted by domain recruitment. Then, two reverse tandem duplications of the ancestral Pif genes resulted in the generation of Pif-d and Pif-e in Pteriomorphia, respectively. In the LCA of Ostreida and Pterioida, Pif-f evolved from a reverse tandem duplication of Pif-e gene, but was lost in Pterioida. Afterwards, the Pif-e gene gained SCR repeat domain by domain shuffling. Finally, VWA domain loss of Pif-f gene resulted in the LamG3 in the LCA of Ostreidae, in which the chalky microstructure appeared in the shells.

**Fig. 6.**
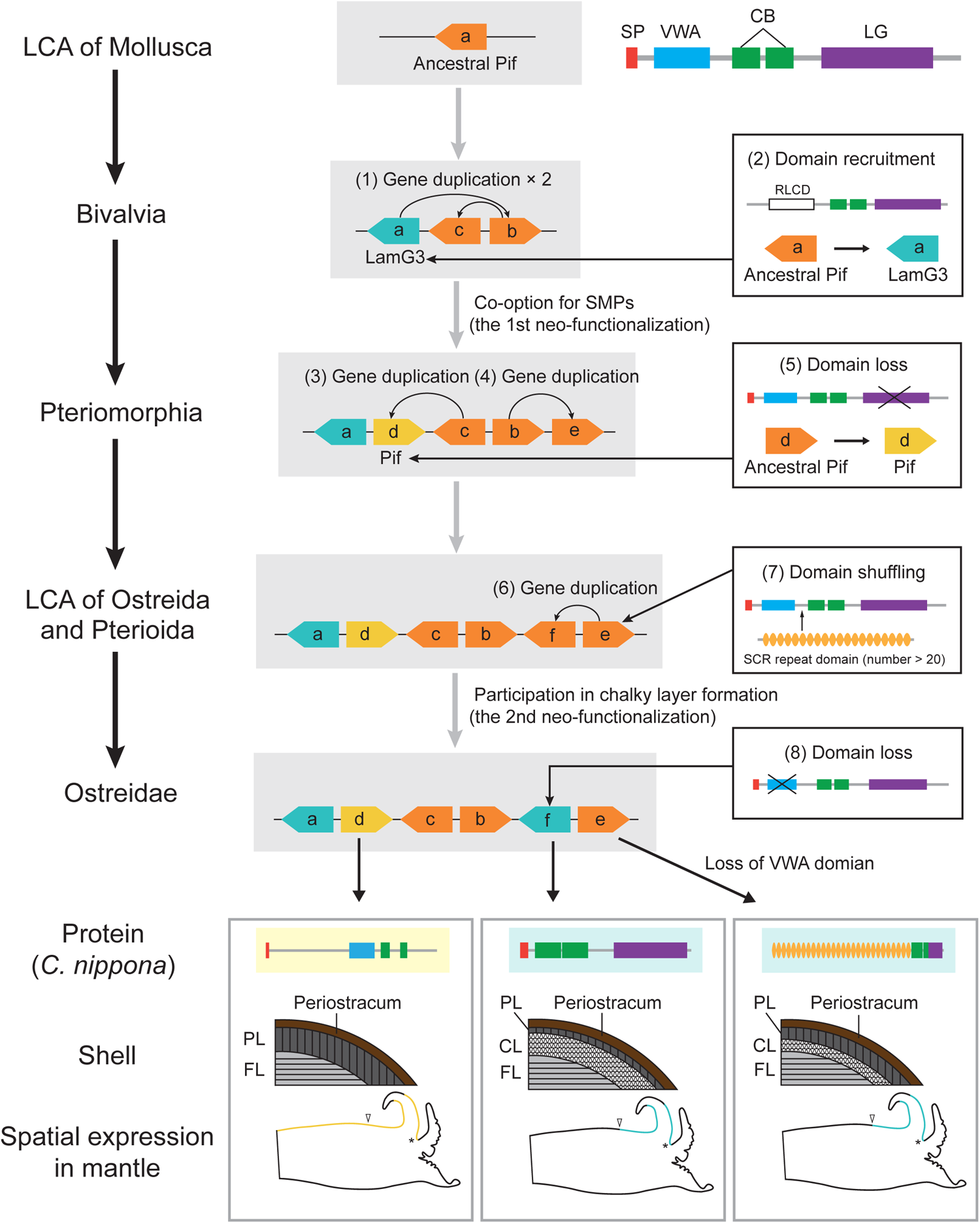
Evolution of the Pif_LamG3_cluster in molluscs. Gray background boxes show the gene cluster of Pif and LamG3. Right squares show the molecular evolution of gene structure. Numbers in brackets indicated the order of molecular evolution events. Asterisks indicate the periostracal groove. Yellow color in the mantle model indicates the expression regions of Pif (CNI_013832-RA), while blue color indicates the expression regions of LamG3 (CNI_014385-RA and CNI_014386-RA). White arrows symbolize the end of the gene expression region. Abbreviations: PL, prismatic layer; FL, foliated layer; CL, chalky layer; CB, chitin-binding domain; VWA, von Willebrand factor type A; LG, concanavalin A-like lectin/glucanase domain; RLCD, repetitive low complexity domain.

The Ostreidae-specific LamG3 evolved as SMP in the chalky layer, and expressed in the dorsal surface of the mantle edge epithelium of *C. nippona*, suggesting its novel function in the oysters. In addition, the Pif-d and Pif-e emerged accompanying the functional evolution that manifested in their high expression level in the mantles of bivalves (supplementary fig. S17) and co-option as SMPs in the *C. nippona* shell (supplementary fig. S15). The Pif-d gene lost the sequence encoding the N-terminal LG domain, and was expressed in the outer epithelial cells of the mantle edge and center (fig 6). The shell layers are ascribed to the compartmentalization of mantle. For example, the outer pallium and mantle edge are responsible for the prismatic layer secretion, whereas the pallial zone and central zone of the mantle secrete the nacreous layer (Marie, et al. 2012). The Pif-d protein was identified in the prismatic and foliated layers of *C. nippona* (fig 6), thereby supporting that the Pif-d gene was involved in the formation of prismatic and foliated layers. The Pif-e gene in *C. nippona* underwent the domain loss of VWA domain, and encoded an SMP involved in the prismatic and chalky layers (fig. 6). The Ostreidae-specific LamG3 gene (Pif-f) and the homolog of Pif-e gene were highly expressed in the dorsal surface of the mantle edge epithelium of *C. nippona*, suggesting that the chalky layer was secreted by the epithelium cells of mantle edge (fig. 6). Strikingly, the three novel genes were all expressed in the inner surface of the outer fold (fig 6), which was responsible for periostracum synthesis (Checa 2000). Periostracum is a tanned organic membrane which is shared by Bivalvia and Gastropoda (de Paula and Silveira 2009). Hence, the conversed function of the ancestral Pif could be associated with the formation of periostracum in Bivalvia. In summary, Pif genes underwent two neo-functionalization events which contributed to shell formation and the diversity of shell microstructures, respectively (fig 6).

Gene duplication from the parental copy usually results in functionally redundant genes which are not stably retained in the genome. In our study, although the duplicated genes were originated commonly from an ancestral Pif, distinct domain architectures enabled them to acquire novel functions involved in the formation of shell microstructures. Such functional evolution of Pif and LamG3 genes may be crucial for their simultaneous retention in the genome. In addition, the spatial expression patterns of these novel genes may be regulated by certain mutation(s) in the gene regulatory region (Shimizu, Takeuchi, et al. 2022). Therefore, understanding evolutionary process of Pif_LamG3_cluster allows us to gain insights into the important role of gene duplication followed by functional diversification in acquiring evolutionary innovation for environmental adaptation.

## Conclusions

In the present study, we report the first high-quality chromosome-level reference genome and comprehensive transcriptomes for the Iwagaki oyster *C. nippona.* Our multi-omic integrative analyses reveal the lineage-specific domains, independent evolution and co-option of genes as the key molecular innovations in the formation and evolution of a novel shell microstructure, chalky calcite. The chalky layer participates in the rapid mineralization of oyster shell and underpins the adaptive innovation of oysters. Comparative analyses of molluscan shell proteomes identified two conserved domains (VWA and CB) in the basic biomineralization toolkit. The ancestral Pif genes originated from the common ancestor of Bilateria were not biomineralization-related. Gene duplication and the dynamic combinations of functional domains enable Pif genes to acquire novel functions for genetic novelties of molluscan shell. Our results supported that key genetic component(s) involved in biomineralization evolved repeatedly from ancestral gene(s) without biomineral function. This study not only provided a framework to deeply understand the origin and evolutionary dynamics of biomineralization in metazoans, but also will facilitate the studies on historical biodiversification events reflected and influenced by biomineralization.

## Methods

### Sample collection and genome sequencing

Multiple wild *C. nippona* individuals were collected from Zhoushan, Donghai Sea, China. The oysters were identified on the basis of both DNA fragments of cytochrome oxidase I (COI) and morphological observation. The samples were dissected, immediately frozen in liquid nitrogen, and stored at −80 □ for further analysis. Genomic DNA was extracted from the adductor muscle of a male individual (674.3 g) by the standard phenol-chloroform method. A PacBio library (15–20 kb) was prepared using the SMRTbell Express Template Prep Kit 2.0. Single-molecule real-time sequencing was carried out on a PacBio Sequel II platform under the CCS mode. Then, the subreads were filtered by minimum length of 50 kb, and the HiFi reads were generated using ccs software (v 4.2.0) (https://github.com/PacificBiosciences/ccs) with the parameters of “min-passes = 3, min-rq = 0.99”. For genome survey, Illumina library was constructed using the DNA from the same oyster, and sequenced by a PE150 strategy on Illumina NovaSeq 6000 system. For Hi-C sequencing, the adductor muscle from the same individual for DNA extraction was fixed with 1 % formaldehyde, and DNA was cross-linked and digested with MboI restriction enzyme. The library was also sequenced on an Illumina NovaSeq 6000 platform with a PE150 method.

### Genome survey, assembly, and scaffolding

Genome survey were performed using k-mer frequency-based method. First, the Illumina reads was trimmed and filtered with fastp (v 0.23.1) (Chen, et al. 2018). The K-mers were then counted using Platanus-allee (v 2.2.2) (Kajitani, et al. 2019). Finally, the output file was used as the input for GenomeScope (v 2.0) (Ranallo-Benavidez, et al. 2020) to estimate the genome size, rate of heterozygosity and abundance of repetitive elements. The initial genome was de novo assembled with hifiasm (v 0.15.1-r328) (Cheng, et al. 2021). The Hi-C sequencing reads were mapped to the contigs by Burrows-Wheeler Aligner (BWA) (v 0.7.17-r1198-dirty) (Li and Durbin 2009). Then, Juicer (v 1.6) (Durand, et al. 2016) was used for the construction of Hi-C contact matrix, and the anchoring was performed with 3D-DNA (v 180419) (Dudchenko, et al. 2017). Finally, the Juicebox Assembly Tools (v 1.11.08) (Dudchenko, et al. 2018) were applied for the manual correction of the connections.

### Quality assessment of genome assembly

To assess the quality of genome assembly, BUSCO (v 5.1.2) (Manni, et al. 2021) analysis was used to verify the completeness of genome. QUAST (v 5.0.2) (Gurevich, et al. 2013) was used to check genome assembly quality with the raw PacBio HiFi reads. In addition, Illumina pair-end reads were mapped back to the assembly with BWA (v 0.7.17-r1198-dirty) (Li and Durbin 2009). Mapping statistics were summarized with samtools software (version 1.15) (Li, et al. 2009).

### Repeat annotation

Repetitive sequences were identified and masked using both homology and de novo predictions. Briefly, RepeatModeler (v 2.0.1) (Flynn, et al. 2020) was used to construct a de novo repeat library. The consensus sequences in de novo repeat library were further combined with molluscan repetitive sequences from both Repbase library (v 20181026) (https://www.girinst.org/repbase/) and Dfam database (v 3.3) (Wheeler, et al. 2013), and then used to run RepeatMasker (v 4.1.2-p1) (Tarailo-Graovac and Chen 2009) on the genome assembly. TE divergence analysis was performed using an R script (https://github.com/ValentinaBoP/Transposable Elements) with the detailed annotation table from the output of RepeatMasker software.

### Gene structure and functional annotation

Transcriptome alignment, de novo prediction, and homology-based methods were combined to predict protein-coding genes in *C. nippona* genome. For transcriptome-based prediction, total RNA was respectively isolated from seven tissues of the same oyster used in genome sequencing (including adductor muscle, digestive gland, gill, male gonad, hemolymph, labial palp, and mantle) as well as female gonad tissue from another wild individual, using TRIzol reagent according to the manufacturer’s instructions. RNA samples from all tissues were pooled (in equal amounts) and used for full-length sequencing on a PacBio Sequel II platform. Raw reads were processed using SMRT Link software (v 9.0) (https://www.pacb.com/support/software-downloads). In addition, RNA-seq short reads generated in our previous research (Gong, et al. 2021) were downloaded from NCBI SRA database (SRR10482020, SRR10482021, SRR10482022, SRR7646736). These downloaded data were pre-processed by fastp (v 0.23.1) (Chen, et al. 2018) and assembled following PASA pipeline (v 2.4.1) (Haas, et al. 2003). High-quality full-length transcripts generated from SMRT Link software and assembled transcripts from PASA were further clustered with cd-hit-est (v 4.8.1) (Li and Godzik 2006). For de novo gene prediction, Augustus (v 3.4.0) (Stanke, et al. 2006) was trained by Braker2 (v 2.1.5) (Brůna, et al. 2021) with short RNA-seq reads. For homologous annotation, protein sequences of *C. gigas*, *C. virginica*, *M. yessoensis*, *Aplysia californica* and *Octopus bimaculoides* were downloaded from NCBI database. Moreover, manually annotated protein sequences (>50aa) of Bivalvia were obtained from the Uniprot/Swiss-Prot database (Release 2022_1). Finally, a high confidence gene set was generated using Maker (v 3.01.03) (Cantarel, et al. 2008) with the trained Augustus predictor, transcript sets and protein sequences from NCBI and Uniprot/Swiss-Prot databases.

Functional annotation of protein-coding genes was carried out by comparing alignments to public databases including NCBI non-redundant (NR), Uniprot/Swiss-Prot, EggNOG (v 5.0) (Huerta-Cepas, et al. 2018), Pfam (Pfam-A v 35.0) (Mistry, et al. 2020), GO categories, and Kyoto Encyclopedia of Genes and Genomes (KEGG) pathways (Kanehisa, et al. 2016). Gene motifs and domains were also identified using InterProScan (v 5.52-86.0) (Jones, et al. 2014). For the annotation of ncRNA, the tRNAscan-SE (v 2.0.7) (Chan, et al. 2021) was employed to predict tRNAs. Screens for rRNAs, miRNAs, and snRNAs were performed using the INFERNAL (v 1.1.2) (Nawrocki and Eddy 2013) against Rfam database (v 14.5) (Griffiths-Jones, et al. 2003).

### Gene family and phylogenetic analyses

The ortholog groups (OGs) of 16 molluscan protein sets were identified using OrthoFinder (v 2.5.2) (Emms and Kelly 2019). Multiple protein sequence alignments were performed with MAFFT (v 7.475) (Katoh and Standley 2013) under default parameters. OGs from selected molluscan taxa were used for subsequent phylogenomic analysis. Phylogenetic tree was constructed based on a total of 1,253 one-to-one single-copy orthologous genes by FastTree 2 (Price, et al. 2010). The MCMCTree (Yang 2007) was used to predict the divergence time among the selected species with nine calibration points (supplementary table S7) obtained from TimeTree database (Kumar, et al. 2017). Expansion and contraction of gene families was estimated using by CAFÉ (v 5) (Mendes, et al. 2020) on the basis of the results from OrthoFinder software (v 2.5.2) and species divergence time. Gene families with P value less than 0.05 were considered as an event of significant expansion or contraction.

To identify tyrosinase, peroxidase, TIMP, VWA, CB, and LG domains, the hmmsearch software was first used to search against the PFAM domain (PF00264.23, PF03098.18, PF00965.20, PF00092.31, PF01607.27, and PF13385.9, respectively) with an E-value threshold of 1e-5. Then, we used InterProScan (v 5.52-86.0) (Jones, et al. 2014) against SMART, Pfam, and SUPERFAMILY databases. Molecular phylogenetic analyses were respectively conducted using tyrosinase, peroxidase, TIMP domain-containing proteins that were identified from 15 protostomian genomes (supplementary table S14) by hmmsearch and InterProScan. Sequence alignment was performed using the program MAFFT (v 7. 475) (Katoh and Standley 2013). The ML phylogenetic trees were constructed using IQ-Tree (v 2.1.4-beta) (Minh, et al. 2020) with 1,000 bootstraps. The final trees were visualized and labeled using iTOL (v 6.7) online (https://itol.embl.de/). For Pif proteins, the phylogenetic tree was built using identified proteins from 16 metazoan genomes (supplementary table S14) following the pipeline described above.

### Synteny analysis

MCscanX in the JCVI toolkit (v 1.1.12) (https://github.com/tanghaibao/jcvi) (Wang, et al. 2012) was used to identify and visualize macro-synteny. We analyzed chromosome collinearity between *C. nippona* and the other three oysters (*C. gigas*, *C. ariakensis*, and *C. virginica*). In addition, syntenic analysis was also performed among *C. nippona*, *M. yessoensis* (Han, et al. 2022), and *P. fucata* (Takeuchi, et al. 2022).

### Transcriptome sequencing and analysis

Farmed *C. nippona* individuals (three-year-old) were collected from the oyster farm of Rushan, Shandong Province, China. For shell regeneration experiment, holes were drilled in the centers of left shells of three oysters. During experiment, shell-damage oysters were cultured in a tank with seawater (seawater temperature of 22 ± 2 □ and salinity of 30 ppt) and fed with *Chlorella* sp. daily. Mantle edge and the central mantle tissue were sampled from left valves of drilled oysters at 10 days post drilling. The other three non-drilled oysters were dissected into adductor muscle, digestive gland, gill, hemolymph, and left mantle tissues (including mantle edge and central mantle).

Total RNA was extracted using TRIzol and further sequenced in PE150 mode on an Illumina NovaSeq 6000 platform to produce ∼ 6 Gb data for each tissue sample. In addition, RNA-seq data from different tissues of *C. gigas*, *Ostrea edulis*, *P. fucata*, *M. yessoensis*, *Mercenaria mercenaria*, and *N. pompilius* were downloaded from NCBI (supplementary table S15). The raw reads of seven species were quality-filtered with fastp (v 0.23.1) (Chen, et al. 2018), and then mapped to their own genomes using HISAT2 (v 2.2.1) (Kim, et al. 2019). For each species, the expression levels of genes were calculated with featureCounts (v 2.0.1) (Liao, et al. 2013) and normalized using Transcripts Per Million mapped reads (TPM) and Transcripts Per Million mapped reads (TMM). The differentially expressed genes (DEGs) for each tissue were identified using edgeR software package (v 3.40.2) (Robinson, et al. 2009). Tissue-specific genes were determined on the basis of their expression levels compared across all tissue types. Specifically, the mantle-specific genes of *C. nippona* were identified with both the mantle edge and central mantle samples against other tissue groups. Only genes which were overexpressed with log_2_(fold change) > 1 and false discovery rate (FDR) < 0.05 against other tissue types were classified as highly expressed genes.

### Real-time PCR validation

To validate our RNA-seq data, quantitative real-time PCR was conducted on selected genes which are highly expressed in the mantle of *C. nippona*, using elongation factor 1-alpha (EF1a) as the internal standard gene. The primers were designed with Primer 6.0 software (supplementary table S16). Real-Time PCR was performed with QuantiNova^TM^ SYBR^®^ Green PCR Kit following the instruction manual of the kit (QIAGEN) on a LightCycler 480 real-time PCR system (Roche). All primer pairs for the PCR amplification were checked by the melting curve method. Three biological replicates for each tissue type were guided. The comparative cycle threshold (Ct) method was applied to quantify the relative expression levels based on the 2^-ΔΔCt^ method (Livak and Schmittgen 2001).

### In situ hybridization

Antisense probes were synthesized using purified PCR products (1 μg per reaction) (supplementary table S16) and DIG RNA labeling Kit (T7) (Roche), following the manufacturer instructions. Probe synthesis reactions were performed at 37 °C for 3 h and then were treated with DNase I (Promega) at 37 °C for 20 min. Synthesized probes were purified using the MEGAclear^TM^ Transcription Clean-Up Kit (Thermo Fisher Scientific) and stored at −80□. Mantle tissues of *C. nippona* were fixed in 4% PFA solution overnight at 4°C. Then, samples were dehydrated with serial methanol (25, 50, 75, and 100%) and stored at −20□.

In situ hybridization of mantles was carried out according to the methods as described previously (Yue, et al. 2021) with slight modifications. Briefly, the fixed mantles were transferred to methanol, cleared in xylene, embedded in paraffin wax, and cut into 5-μm-thick sections on a Leica RM 2016 rotary microtome (Leica). After a series of deparaffinization, hydration, digestion, prehybridization, hybridization (final concentration of RNA probes: 1 ng/μl), and antibody incubation (with a 1:3000 dilution of antDIG-AP antibody in the blocking buffer), sections were incubated with 2% NBT/BCIP solution (Roche) in darkness at 4 °C overnight. Finally, pictures were taken under an Olympus BX53 microscope coupled with a DP80 camera (Olympus).

### Scanning electron microscopy

To characterize crystal structures, the *C. nippona* shells were fractured and carefully collected with a dissecting knife under an anatomical lens. After a 5 min ultrasonic cleaning, the shells were dried and sputter-coated with a thin layer of gold nanoparticles. Then, the surfaces and vertical sections of shells were scanned using the VEGA3 TESCAN scanning electron microscope.

### LC-MS/MS analysis

Fresh shells of six *C. nippona* individuals were incubated in 1% sodium hypochlorite (NaOCl) for 24 h and mechanically washed in the Milli-Q water to remove remaining tissues, superficial epibionts, and periostracum. The outer prismatic, inner foliated and chalky layers were identified by their color, and carefully separated using a dissecting knife. Separated shell layers were cleaned with a 5 min ultrasound treatment in the Milli-Q water and then air-dried at room temperature (RT).

The cleaned shell layers were roughly crushed into fine powder and treated with SDT-lysis buffer (4% SDS, 100mM DDT, 100mM Tris-HCl) in a boiling water bath for 5 min. After cooling to RT, the supernatant was collected by a short centrifugation, then mixed with UA buffer (8M Urea, 150mM Tris-HCl, pH 8.0). The mixture was ultra-filtered on 10 kDa cut-off membrane, and alkylation was performed with 50mM iodoacetamide in UA buffer for 30min at RT in the dark. After washing with UA buffer and NH_4_HCO_3_ solution sequentially, samples were digested with trypsin solution (6µg trypsin in 40µl NH_4_HCO_3_ buffer) at 37 °C for 16 h, desalted via C18 Stage Tips and dried off in a vacuum concentrator. The dried peptides were then reconstituted in 0.1% formic acid for analysis by a Q-Exactive Plus mass spectrometer coupled to an EASY-nLC 1200 system (Thermo Fisher Scientific).

Peptide fragments were analyzed against the predicted gene models of *C. nippona* using the intensity-based absolute quantification (iBAQ) method in MaxQuant (v 1.6.17.0) (Tyanova, et al. 2016). Minor and major proteins were discerned following the procedures as previously described (Mann and Edsinger 2014). In addition, amino acid sequences of minor proteins were searched against SMP database (https://doi.org/10/cz2w) (Yarra, et al. 2021) and shell proteome of *C. gigas* (Zhao, et al. 2018), using BLASTP (v 2.11.0) with an E-value of 1e-100 and sequence identity of 80%. Major proteins and the best matches of minor proteins were identified as SMPs of *C. nippona* in this study. Furthermore, SMPs of the other six molluscs including *C. gigas* (Zhao, et al. 2018), *Atrina pectinata* (Shimizu, Negishi, et al. 2022), *Tridacna crocea* (Takeuchi, et al. 2021), *P. fucata* (Zhao, et al. 2018), *L. gigantea* (Marie, et al. 2013), and *N. pompilius* (Zhang, et al. 2021) were download and used for comparative analysis of shell proteomes. Functional domain annotations of SMPs were performed by searching against various databases, including SMART, CDD, Pfam, PROSITE patterns, PROSITE profiles, and SUPERFAMILY, using InterProScan (v 5.52–86.0) (Jones, et al. 2014).

## Supplementary Information

Supplementary data are available in the supplementary file (Supplementary Material).

## Authors’ contributions

Y. B., S. L., and Q. L. conceived the idea and project. Y. B. and Y. H. collected the samples. Y. B. performed genomic, transcriptomic and shell proteomic analyses. H. Y., L. K., and C. X. contributed materials and reagents. Y. B. performed the SEM, RT-PCR, and ISH experiments. Y. B., S. L., and Q. L. drafted and revised the article. All authors read and approved the final manuscript.

## Availability of data and materials

All raw genome and transcriptome sequencing data used for genome assembly and annotation have been deposited at NCBI under the BioProject accession PRJNA947686. The RNA-seq data from various tissue transcriptomes of *Crassostrea nippona* (SRR23950953-SRR23950959, SRR23950967, SRR23950970-SRR23950979) and shell-repair experiment (SRR23950960-SRR23950965) were deposited in NCBI SRA database under PRJNA947922 project. The shell proteomic data, genome assembly and annotation have been deposited on Figshare (doi: 10.6084/m9.figshare.22336354). The code commands used in this study are available in the supplementary file (Supplementary scritps.txt).

## Declarations

### Ethics approval and consent to participate

Not applicable.

### Consent for publication

Not applicable.

