## supplementary fig. S1-S17 for "Neo-functionalization and co-option of Pif genes facilitate the evolution of a novel shell microstructure in oysters"

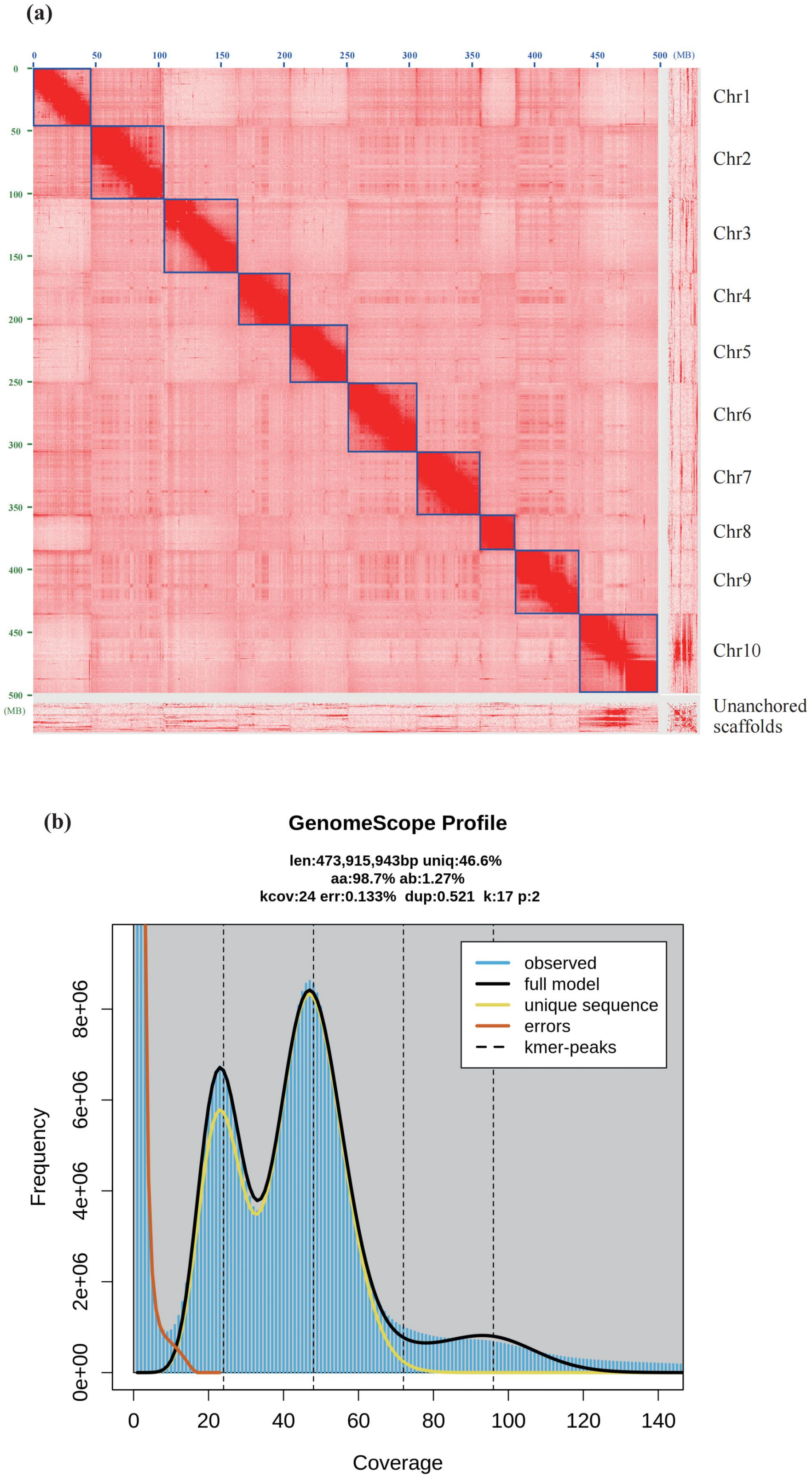


**Figure S1. Genome assembly of *Crassostrea nippona*.** (a) The Hi-C heatmap of genome assembly. The righ axis represents the chromosome number. (b) Estimated genome size of *C. nippona* based on K-mer analysis.

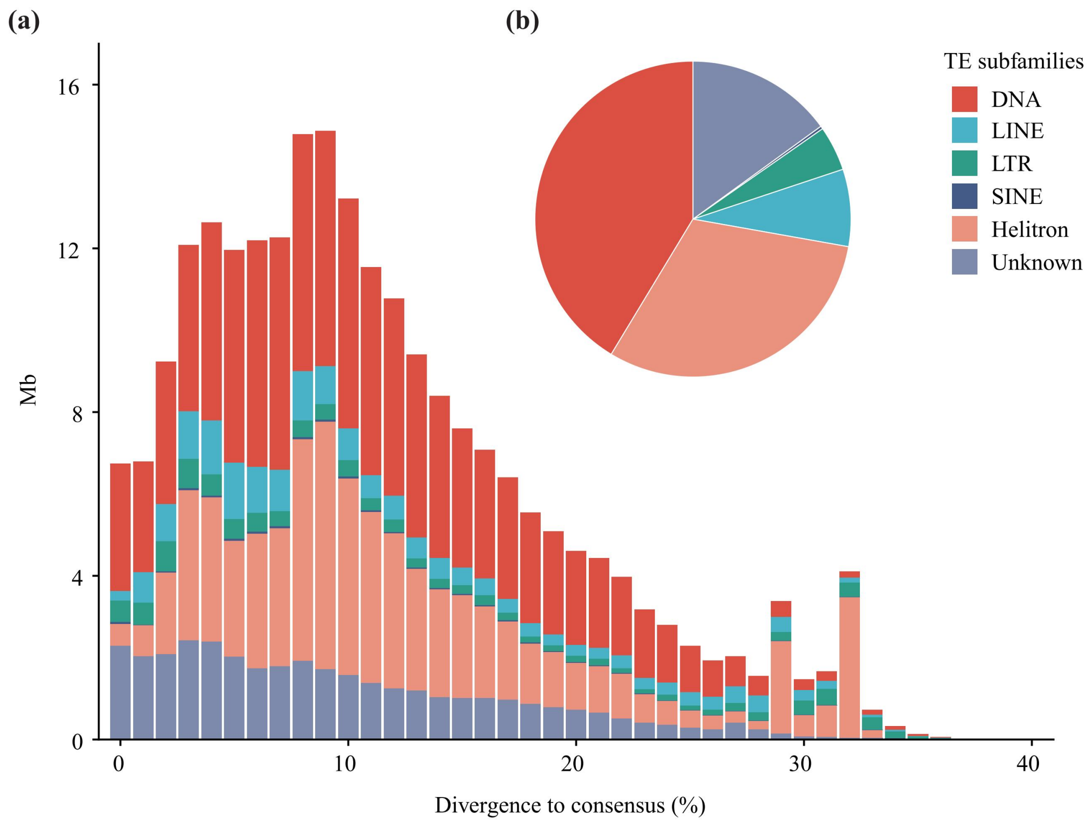


**Figure S2. Distribution of TEs in the *Crassostrea nippona* genome.** (a) History of TE accumulation in the *C. nippona* genome. Historical TE divergence was calculated by the Kimura distance-based copy divergence analysis. (b) Proportions of Helitrons, DNA transposons, LTR, LINE and SINE retrotransposons in the *C. nippona* genome.

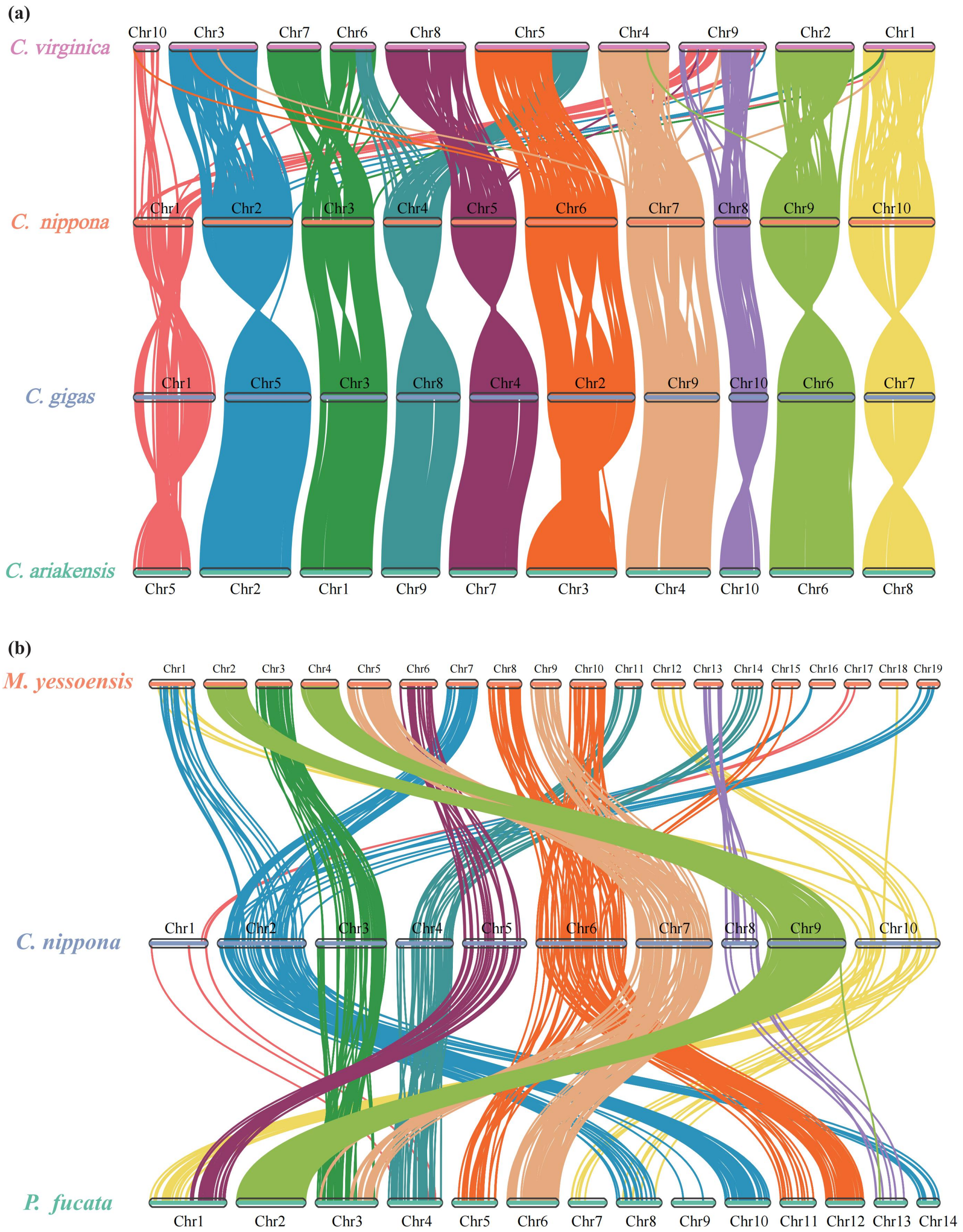


**Figure S3. Genomic synteny between *Crassostrea nippona* and other molluscs.** (a) Macro-synteny among four Crassostrea species. (b) Macro-synteny comparisons across *C. nippona*, *Mizuhopecten yessoensis*, and *Pinctada fucata*.

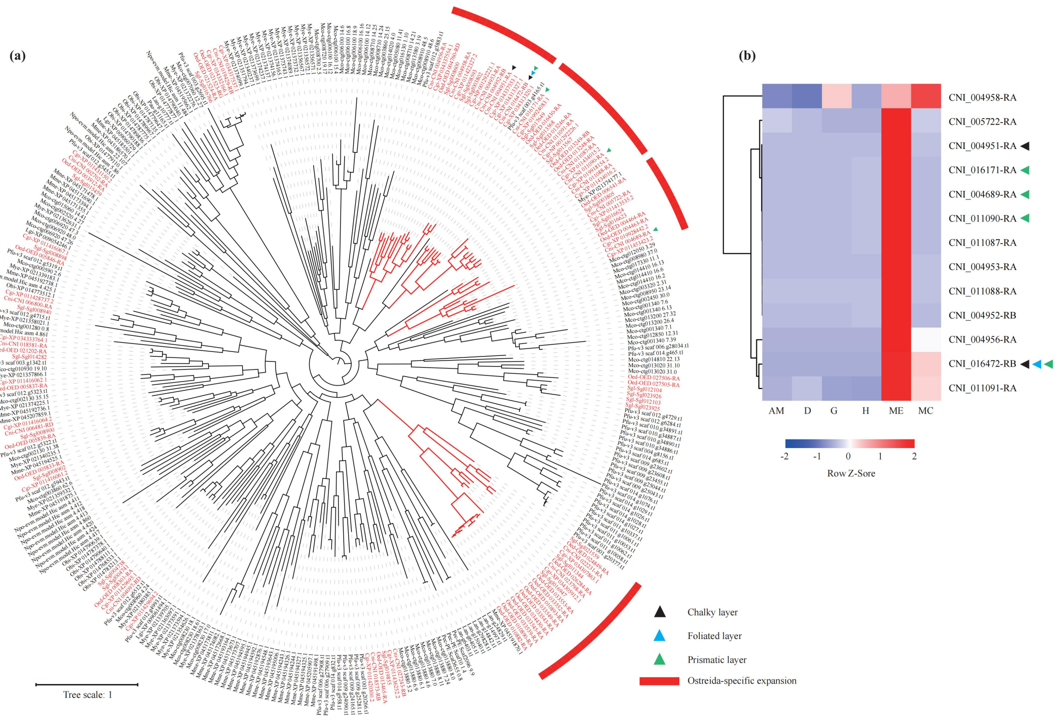


**Figure S4. Expansion of tyrosinase gene family in Protostomia.** (a) Phylogeny of protostomian tyrosinases demonstrates oyster-specific expansions. (b) Expression pattern of oyster-specific expanded tyrosinase genes in *Crassostrea nippona*.

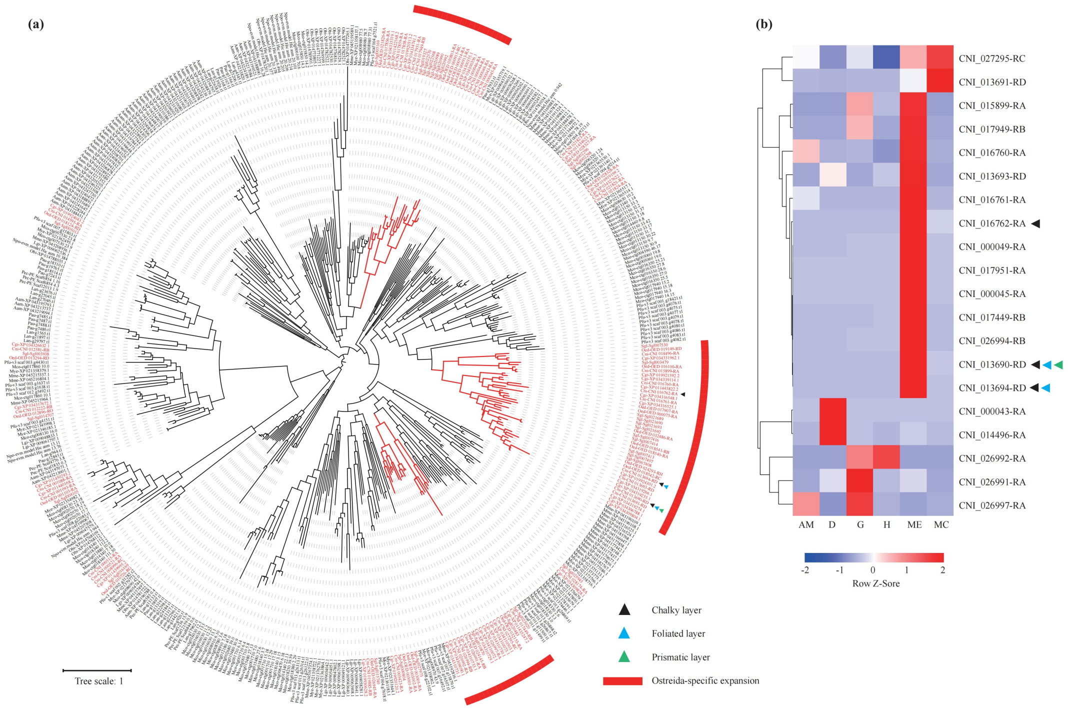


**Figure S5. Expansion of peroxidase gene family in Protostomia.** (a) Phylogeny of protostomian peroxidases demonstrates oyster-specific expansions. (b) Expression pattern of oyster-specific expanded peroxidase genes in *Crassostrea nippona*.

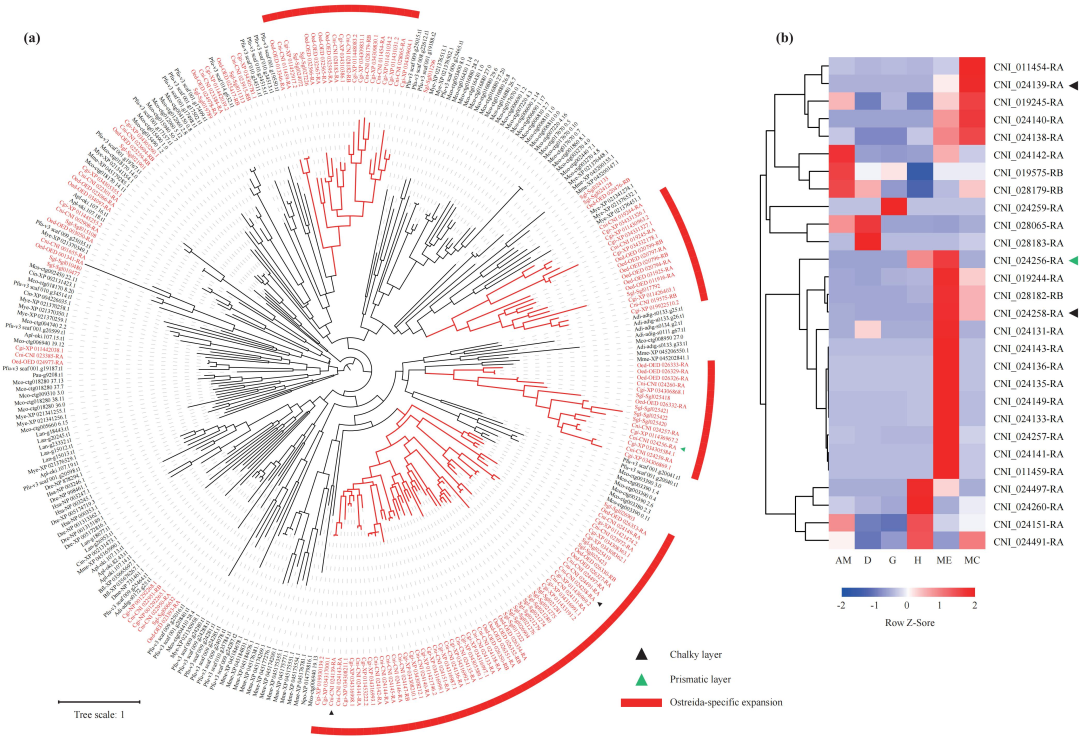


**Figure S6. Expansion of tissue inhibitor of metalloproteinase (TIMP) gene family in Protostomia.** (a) Phylogeny of protostomian TIMPs demonstrates oyster-specific expansions. (b) Expression pattern of oyster-specific expanded TIMP genes in *Crassostrea nippona.*

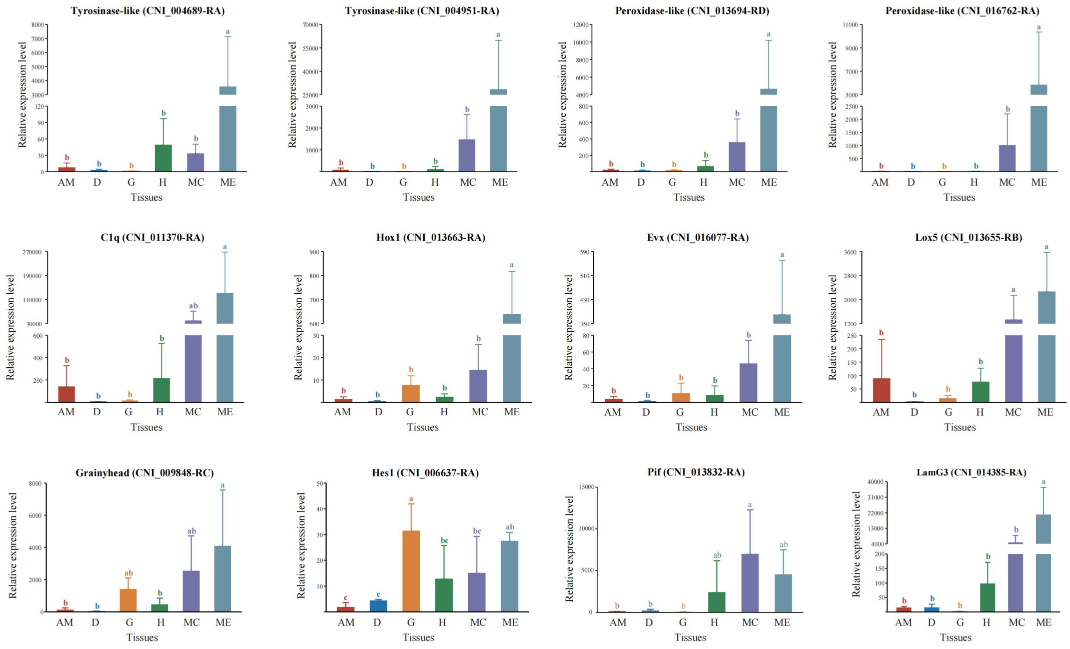


**Figure S7. Real-time PCR results showing gene expression patterns among tissues of *Crassostrea nippona* (n = 3 biologically independent individuals).** Statistical significance ($P < 0.05$) shown by different letters was determined using the LSD test in the R package agricolae (v 1.3-5). Abbreviations as fig. S7.

(a)

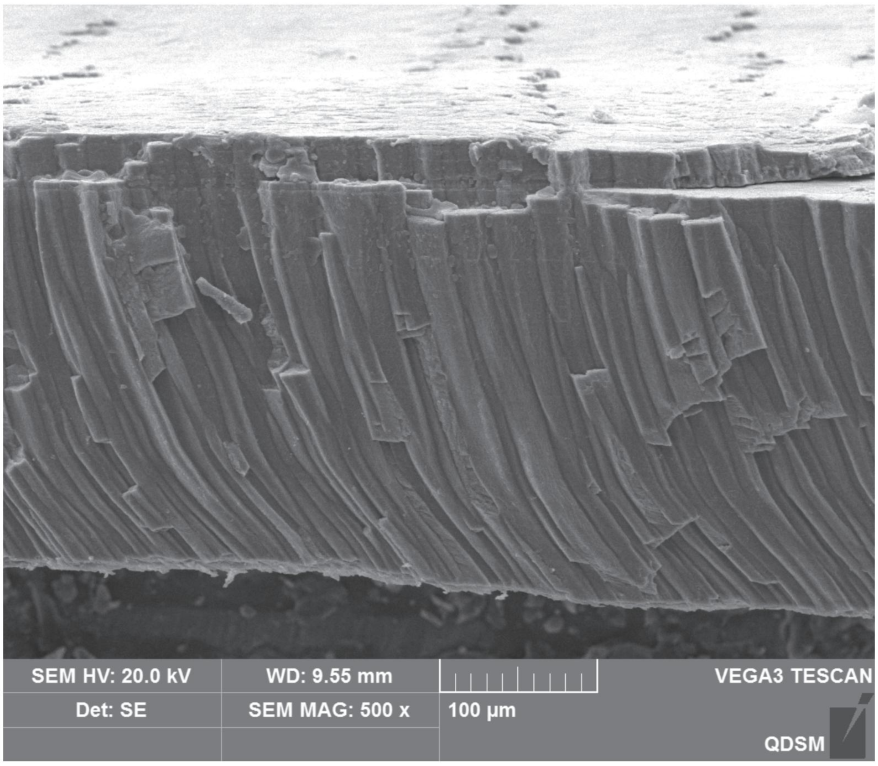


(b)

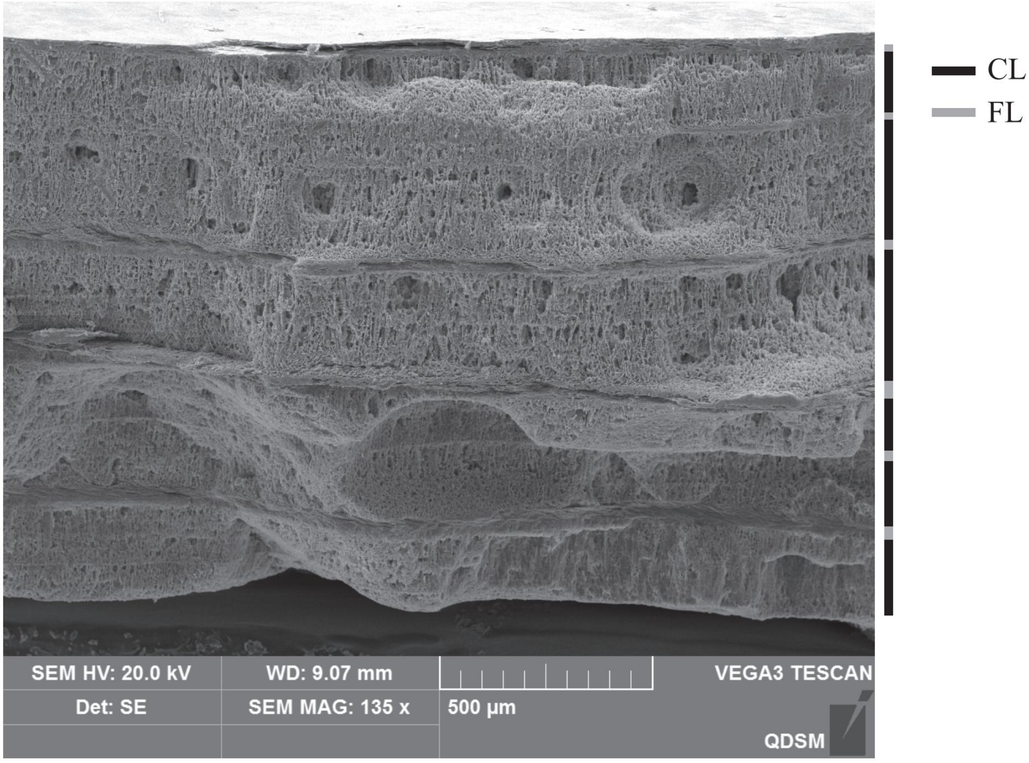


**Figure S8. Ultrastructure of the *Crassostrea nippona* shell.** (a) Outer layer of prisms. (b) Inner multi-layered structures. Abbreviations: FL, foliated layer; CL, chalky layer.

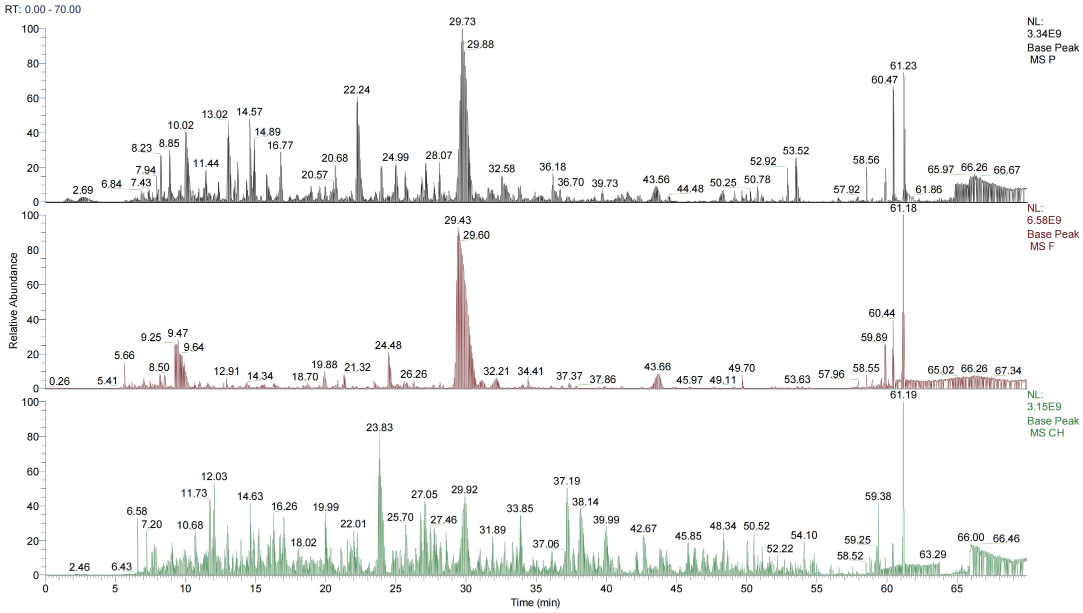


**Figure S9. Base peak chromatogram of three types of protein sample of the *Crassostrea nippona* shell.** Black color indicates protein from the prismatic layer; Red color: foliated layer; Green color: chalky layer.

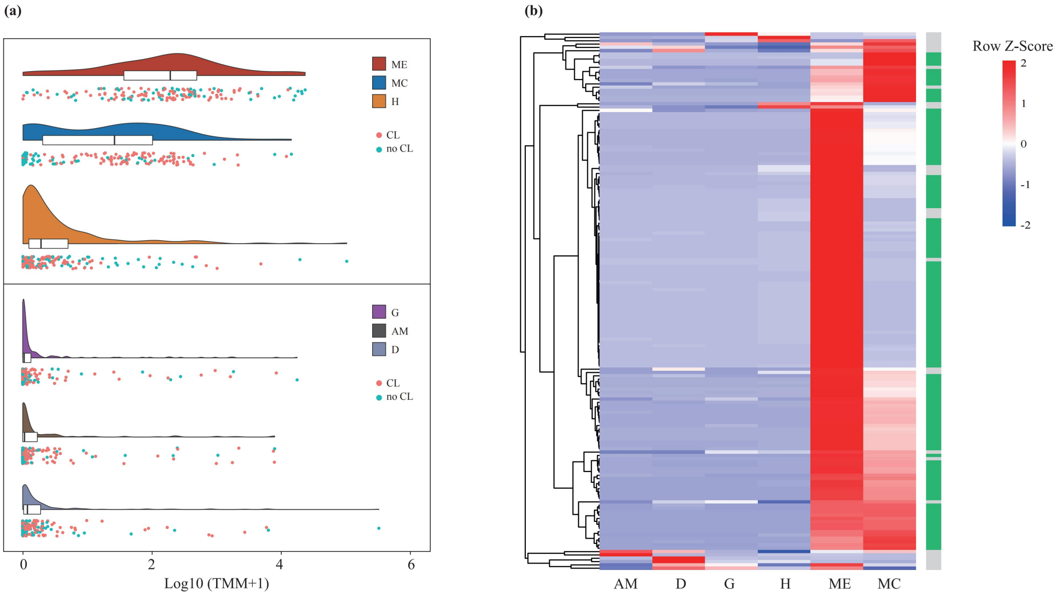


**Figure S10. Expression patterns of genes encoding shell matrix proteins (SMPs) in six types of tissues of *Crassostrea nippona*.** Abbreviations: AM, adductor muscle; D, digestive gland; G, gill; H, hemolymph; ME, mantle edge; MC, central mantle. (a) Distribution of expression levels of genes encoding SMPs in six types of tissues. Red dots indicate SMPs involved in the chalky layer. Blue dots indicate SMPs which are not identified in the chalky layer. Abbreviations: TMM, Transcripts Per Million mapped reads. (b) Tissue-specific expression of SMPs in *C. nippona*. Heatmap shows the normalized expression profiles of SMPs in different tissues. Mantle-specific genes encoding SMPs are marked with green color on the right.

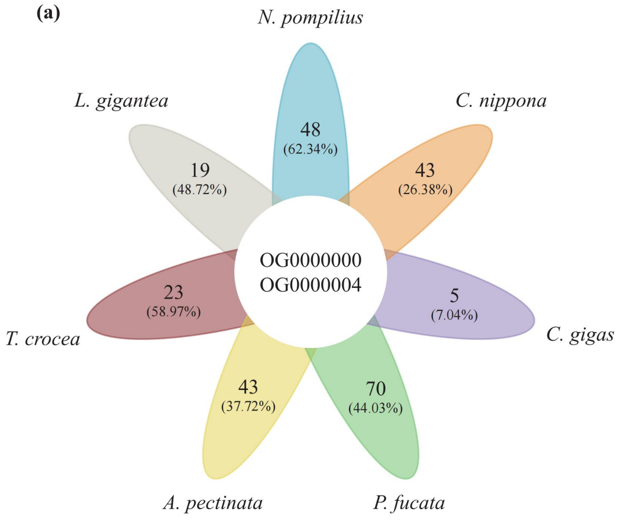


**Figure S11. OrthoFinder analysis of shell proteomes from seven molluscs.** (a) Flower plot t comparing orthologous groups and species-specific SMPs among seven species. Numbers show the species-specific SMPs of each species, while percentages in brackets indicate the proportion of species-specific SMPs in each species. (b) Number of *C. nippona*-specific SMPs identified from the prismatic (PL), foliated (FL), and chalky layers (CL).

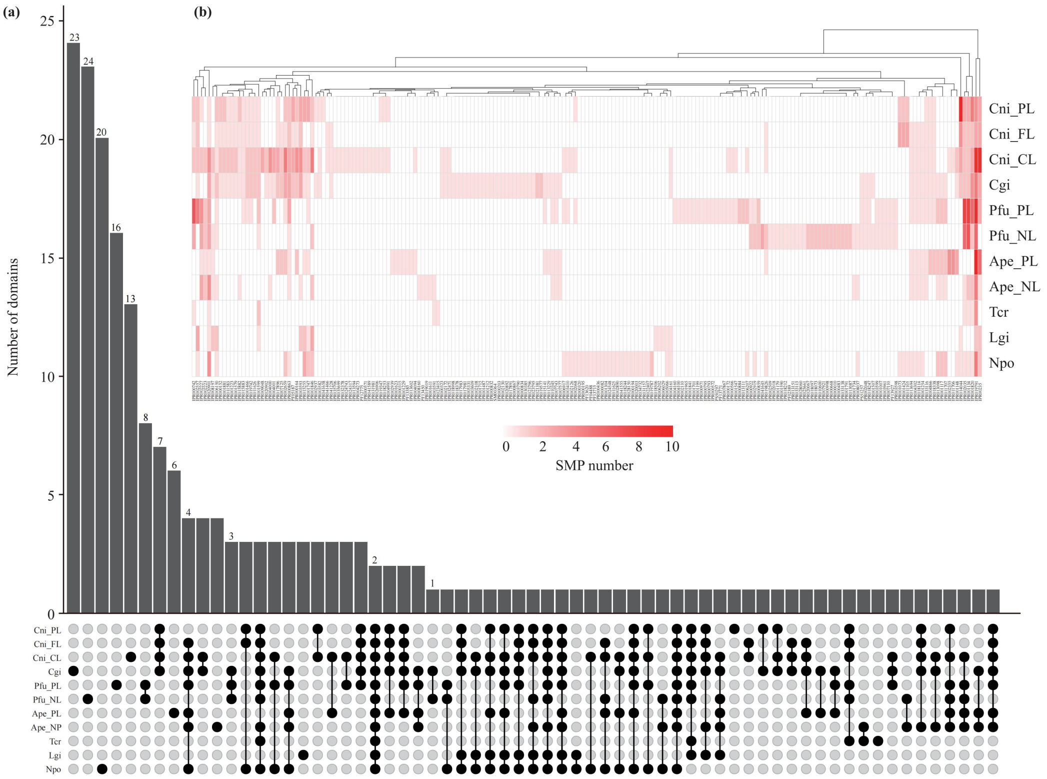


**Figure S12. Protein-domain analysis of shell proteomes from seven molluscs.** (a) Complete result of comparing the domains identified from the shell proteomes. (b) Distribution of SMP domains in different shell microstructures of seven species. Abbreviations as fig. S7, except: Ape, *A. pectinate*; Cgi, *C. gigas*; Cni, *C. nippona*; Pfu, *P. fucata*; Tcr, *T. crocea*; Lgi, *L. gigantea*; Npo, *N. pompilius*.

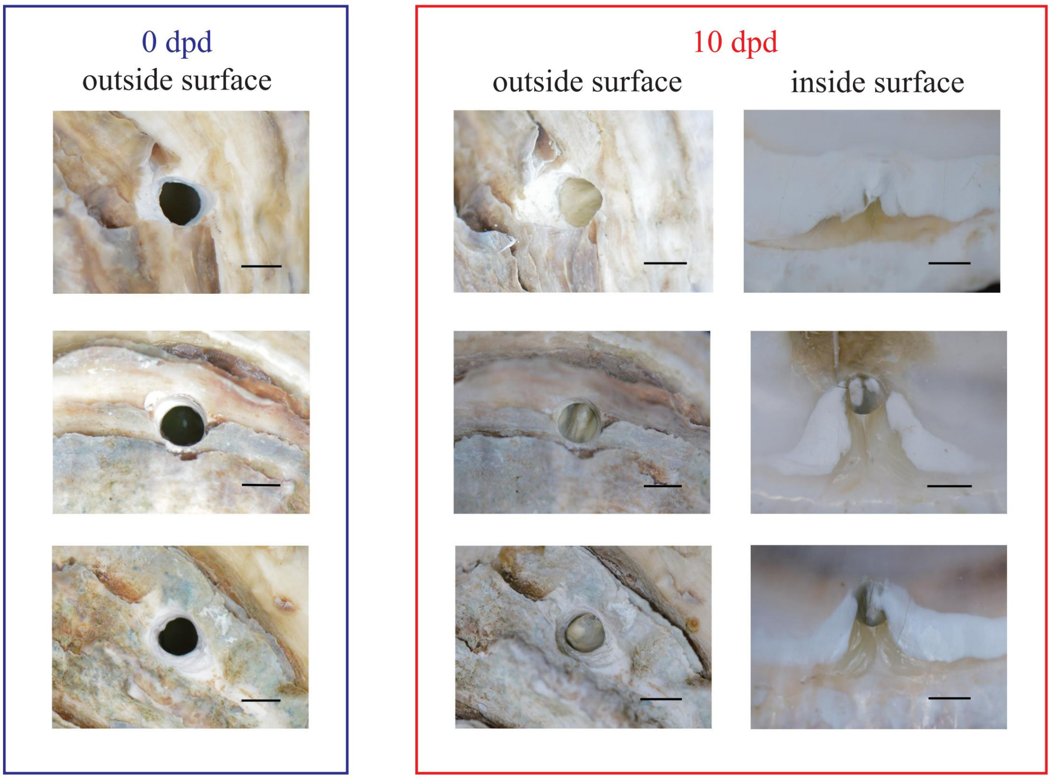


**Figure S13. Observation of the shell repair process of *Crassostrea nippona*.** Abbreviations: dpd, days post shell-drilling Scale bar, 5 mm.

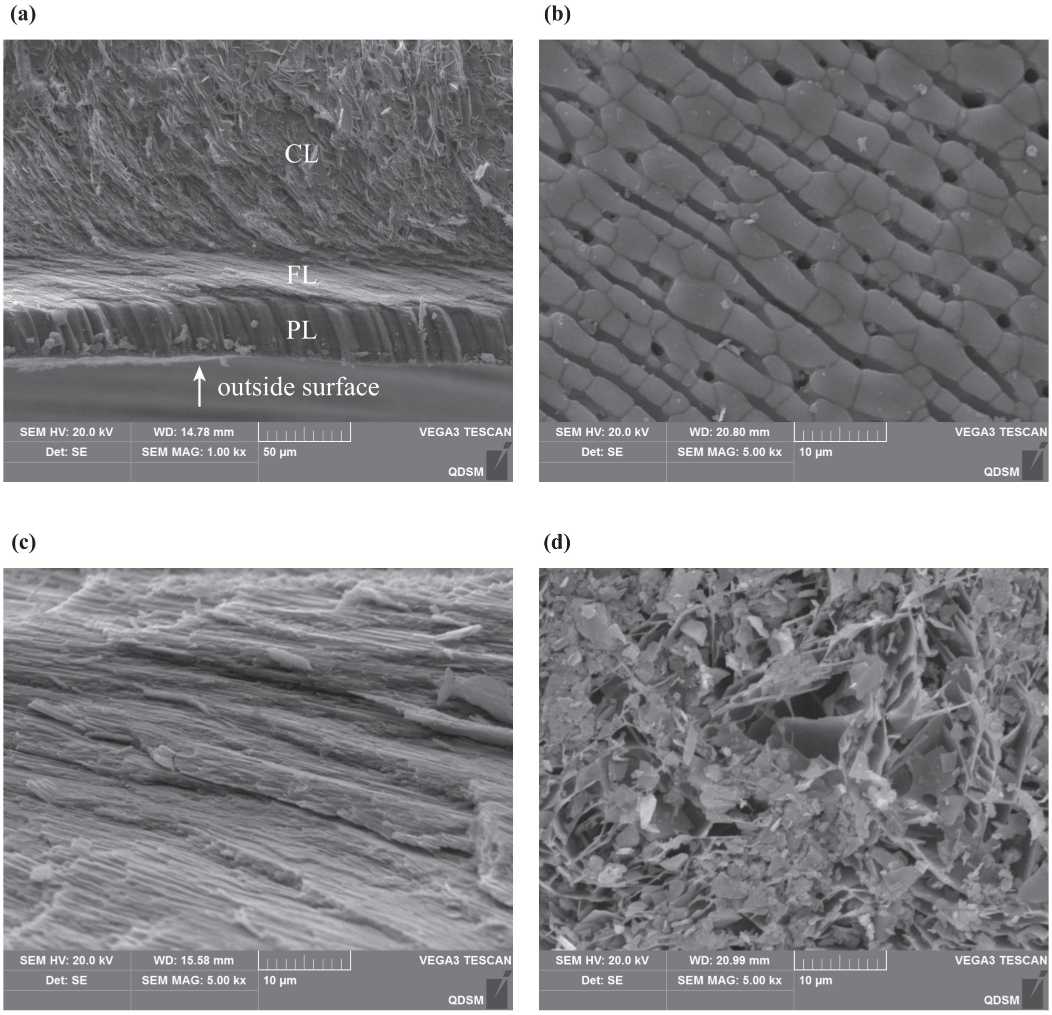


**Figure S14. SEM images representative of the ultrastructure of repaired shell of *Crassostrea nippona*.** (a) Cross section of the whole repaired shell. (b) Repaired surface of PL. (b) Cross section of FL. (d) Cross section of CL. Abbreviations as fig. S8.

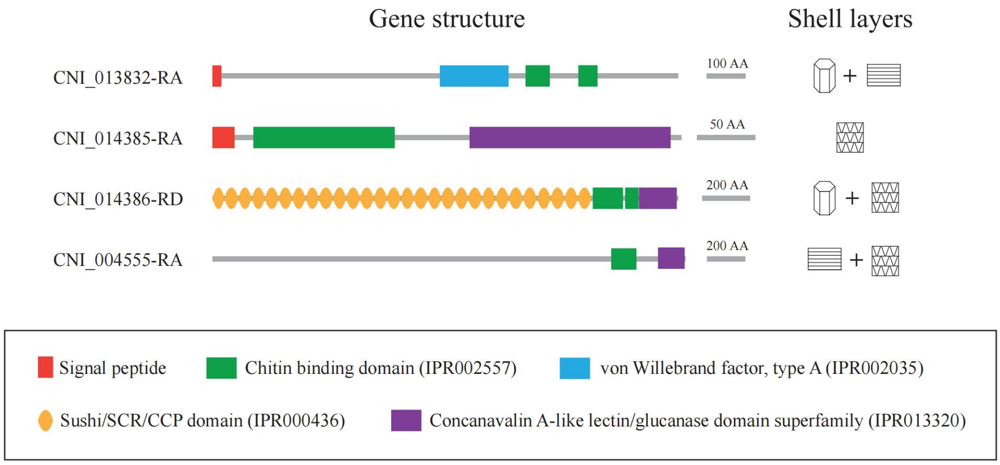


**Figure S15. Cartoon representation indicating domains in Pif and LamG3 proteins identified as SMPs of *Crassostrea nippona*.** Microstructure models as in fig 2d.

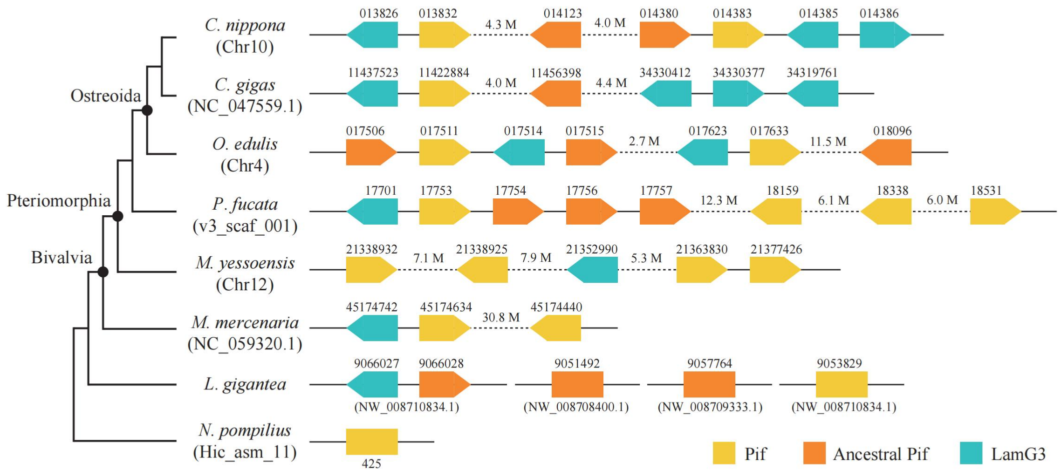


**Figure S16. Genomic arrangement of Pif, ancestral Pif, and LamG3 genes in mollusks.** Arrows indicate the direction of the transcripts. Dashed lines represent the long gaps in the genome.

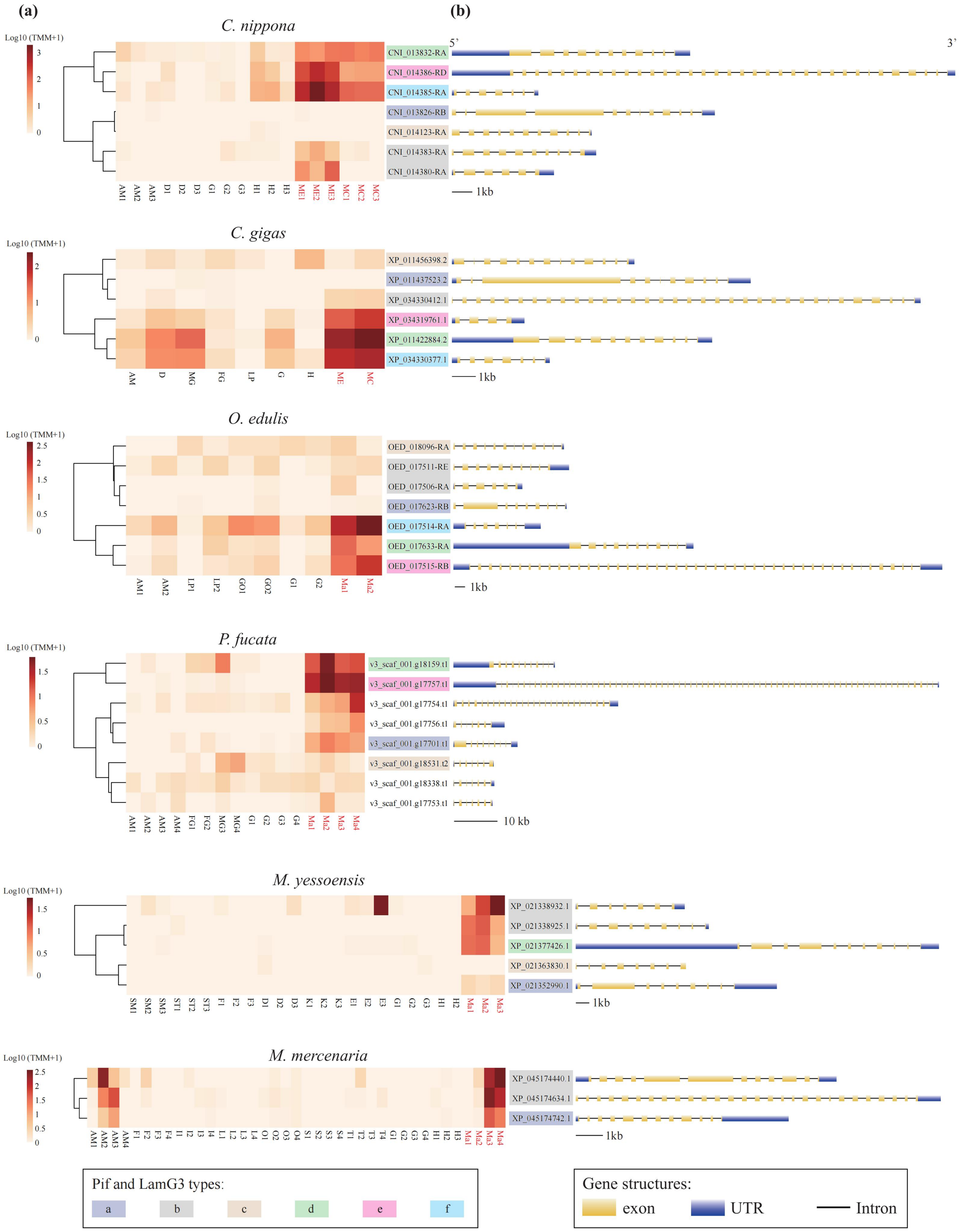

(a)
(b)
C. nippona
Log10 (TMM+1)
5'
3'
CNI_013832-RA
CNI_014386-RD
CNI_014385-RA
CNI_013826-RB
CNI_014123-RA
CNI_014383-RA
CNI_014380-RA
1kb
C. gigas
XP_011456398.2
XP_011437523.2
XP_034330412.1
XP_034319761.1
XP_011422884.2
XP_034330377.1
O. edulis
OED_018096-RA
OED_017511-RE
OED_017506-RA
OED_017623-RB
OED_017514-RA
OED_017633-RA
OED_017515-RB
P. fucata
v3_scaf_001.g18159.t1
v3_scaf_001.g17757.t1
v3_scaf_001.g17754.t1
v3_scaf_001.g17756.t1
v3_scaf_001.g17701.t1
v3_scaf_001.g18531.t2
v3_scaf_001.g18338.t1
v3_scaf_001.g17753.t1
10 kb
M. yessoensis
XP_021338932.1
XP_021338925.1
XP_021377426.1
XP_021363830.1
XP_021352990.1
M. mercenaria
XP_045174440.1
XP_045174634.1
XP_045174742.1
Pif and LamG3 types:
a
b
c
d
e
f
Gene structures:
exon
UTR
Intron

**Figure S17. Tissue expression patterns and gene structures of Pif_LamG3_cluster members in bivalves.** (a) Expression heatmaps of Pif_LamG3_cluster members in different tissues of bivalves. Mantle tissues are marked by red font. The number after tissue name represents the biological duplicate. Abbreviations: AM, adductor muscle; D, digestive gland; E, eyes; F, foot; FG, female gonad; G, gill; GO, gonad; H, hemolymph; I, intestine; K, kidneys; L, liver; LP, labial palps; Ma, mantle; MC, mantle center; ME, mantle edge; MG, male gonad; O, ovary; S, stomach; SM, striated muscle (adductor muscle); ST, striated muscle (adductor muscle); T, testis. (b) Gene structure of Pif_LamG3_cluster members.
